# Loss of *Usp22* enhances histone H2B monoubiquitination and stimulates intracellular and systemic interferon immunity

**DOI:** 10.1101/2021.04.09.439190

**Authors:** Nikolaus Dietlein, Xi Wang, Esther Rodríguez Correa, Beyza Erbil, Daniel B. Lipka, Yvonne Begus-Nahrmann, Robyn Laura Kosinsky, Steven A. Johnsen, Thomas H. Winkler, Hans-Reimer Rodewald

## Abstract

Interferons protect from virus infections by inducing hundreds of interferon-stimulated genes (ISG) which orchestrate anti-viral adaptive and innate immunity. Upon viral infection or type I interferon (IFN) stimulation of cell lines, a histone modification, monoubiquitinated histone 2B (H2Bub1), increases at ISG loci, raising the possibility that a specific chromatin state can broadly stimulate IFN immunity in vivo. Here we show that, in the absence of virus infection or elevated interferon levels, mice lacking the relevant deubiquitinase, *Usp22*, in immune cells have elevated H2Bub1 levels. Hypermonoubiquitinated H2B is physically associated with dozens of ISG loci, and expression of large numbers of ISG is upregulated. This epigenetic state promotes intracellular and systemic immune phenotypes akin to adaptive and innate interferon immunity, and thereby identifies *Usp22* as a negative regulator of interferon immunity.

## Introduction

Interferons are cytokines produced upon viral and bacterial infections which exert their function through the induction of several hundred ISG. On the cellular level, ISG expression confers an anti-viral state by a plethora of mechanisms, including inhibition of viral entry and replication (reviewed in Schneider et al., 2014). Organismal interferon-mediated effects include HSC activation (Baldridge et al., 2010; Essers et al., 2009), increased myelopoiesis (Buechler et al., 2013), impaired B cell development (Lin et al., 1998), spontaneous T cell activation as well as increased germinal center formation, enhanced plasma cell differentiation and increased immunoglobulin production (Jego et al., 2003) (reviewed in Ivashkiv and Donlin, 2013). While it is well established that the activation of STAT transcription factors downstream of IFN receptor engagement drives the expression of ISG and thereby mediates the effects of IFN (reviewed in Schneider et al., 2014), it is unclear whether there is a unifying chromatin state underlying such diverse responses across the immune system.

Monoubiquitinated H2B (H2Bub1) is an activating chromatin mark found in the transcribed regions of highly expressed genes (Minsky et al., 2008). In mammalian cells, H2B monoubiquitination is catalyzed by the RNF20/RNF40 E3 ubiquitin ligase complex (Zhu et al., 2005). While perturbation of H2Bub1 levels only affects a minor fraction of the transcriptome (Shema et al., 2008; Xie et al., 2017), especially inducibly transcribed genes are sensitive to impaired H2B monoubiquitination (Shukla and Bhaumik, 2007; Prenzel et al., 2011; Shema et al., 2008). Upon viral infection or type I interferon stimulation of cell lines, H2Bub1 levels increase (Fonseca et al., 2012). USP22 is the enzymatically active component of the SAGA deubiquitinase module, and acts as an H2Bub1 deubiquitinase (Zhang et al., 2008a; Zhao et al., 2008). Collectively, these findings raise the intriguing possibility that increased levels of H2B monoubiquitination induce ISG expression and stimulate interferon phenotypes in vivo even in the absence of an external interferon signal or a virus infection. Here, we tested this idea by generation followed by cellular and molecular analysis of mice lacking *Usp22* in the immune system, in which H2Bub1 levels are enhanced via loss of the deubiquitinase.

## Results

### Consequences of *Usp22* gene deletion on H2B monoubiquitination, and expression of interferon-stimulated genes

We first examined whether *Usp22* regulates H2Bub1 levels in hematopoietic cells in vivo. Given that global loss of *Usp22* is embryonically lethal (Lin et al., 2012), we generated *Usp22^fl/fl^ Vav1-Cre* mice with pan-hematopoietic *Usp22* deletion (termed *Usp22* KO) (Fig. 1A). Single cell genotyping revealed efficient deletion of the *Usp22* gene across the immune system (Fig. 1B). Consequently, *Usp22* mRNA expression was largely abrogated (Fig. 1C). At histones, levels of monoubiquitinated H2B were determined by western blotting, using total H2B as loading control. Across all tested hematopoietic cell types (for cell types see figure legends) H2Bub1 levels were markedly enhanced in the absence of *Usp22* (Fig. 1D; see Materials and methods). Hence, *Usp22* is a non-redundant H2Bub1 deubiquitinase in hematopoietic cells.

**Figure 1.**
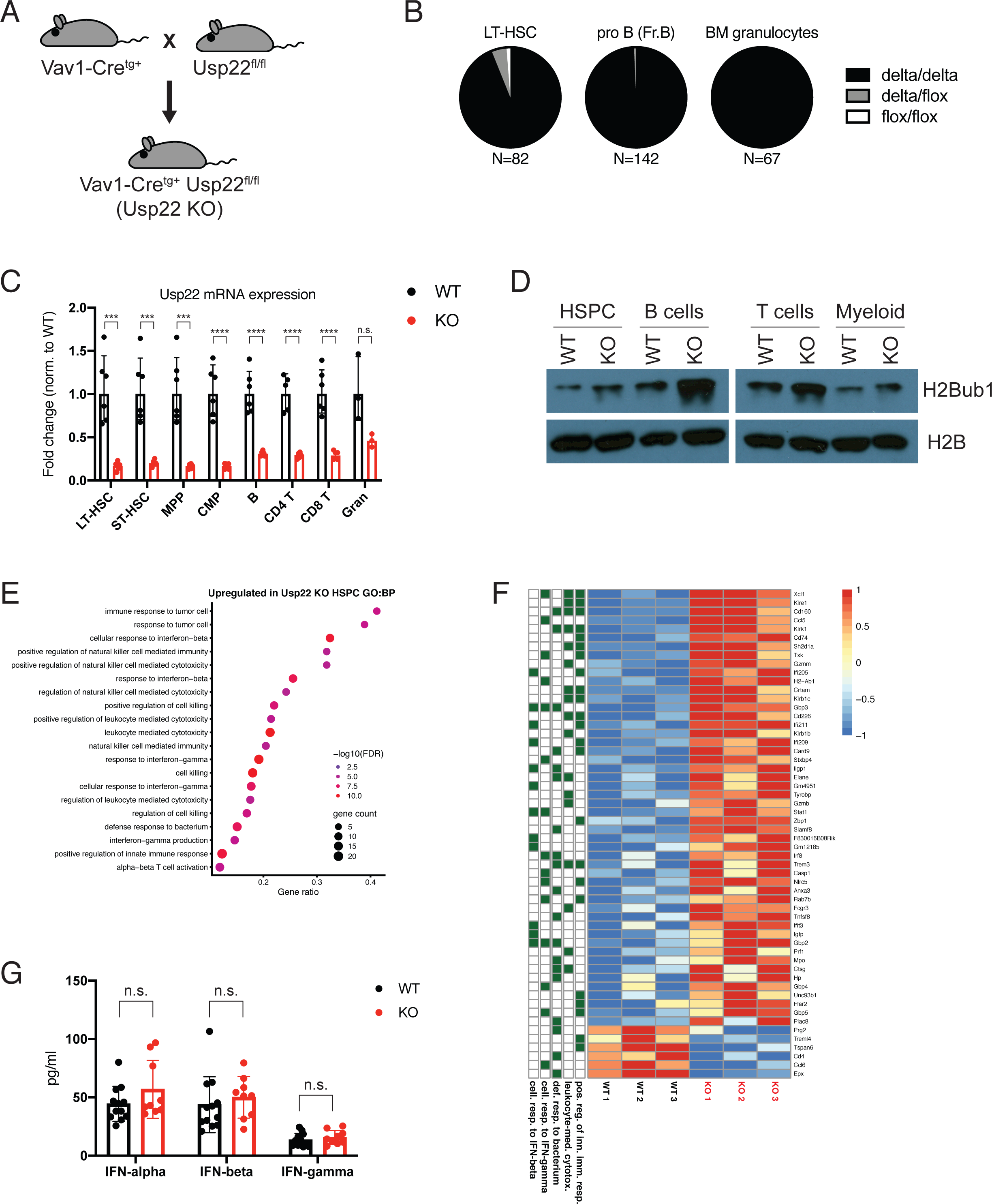
*Usp22* deficiency drives inflammatory gene expression in HSPC. (**A**) Transgenic Vav1-Cre mice were crossed with mice carrying a floxed *Usp22* allele to obtain mice with pan-hematopoietic *Usp22* deficiency (*Usp22* KO). (**B**) *Usp22* genotype of individual cells within the indicated cell populations isolated from *Usp22* KO mice determined by single-cell PCR (n=67-142 cells) (delta/delta = KO). (**C**) Quantitative reverse transcription (qRT-)PCR analysis for *Usp22* mRNA expression in long-term hematopoietic stem cells (LT-HSC), short-term hematopoietic stem cells (ST-HSC), multipotent progenitors (MPP) and common myeloid progenitors (CMP) from bone marrow, and B cells, CD4^+^ T cells, CD8a^+^ T cells and granulocytes from the spleens of wild type and *Usp22* KO mice (n=3-6). (**D**) Western Blot analysis for H2B and H2Bub1 in bone marrow HSPC, and splenic B (CD19^+^B220^+^), T (both CD4^+^ and CD8a^+^ cells) and myeloid cells (CD11b^+^) from wild type and *Usp22* KO mice (n=8 mice per genotype). (**E**) GO term analysis of DEG upregulated in *Usp22* KO versus wild type HSPC. (**F**) Heatmap showing the expression of genes differentially expressed between wild type and *Usp22* KO HSPC within selected GO terms identified in (**E**). (**G**) Interferon serum levels in wild type and *Usp22* KO mice (n=9-12). Geometric mean ± geometric s.d. (**C**), mean ± s.d. (**G**). * p<0.05; ** p<0.01; *** p<0.001; **** p<0.0001; n.s., not significant by unpaired, two-tailed Student’s t-test.

Gene expression analysis in wild type and *Usp22 KO* cells by RNA sequencing identified 380 differentially expressed genes (DEG) (212 upregulated and 168 downregulated in *Usp22* KO versus wild type) (Fig. S1A), hence only a minor fraction of the transcriptome (Shema et al., 2008; Xie et al., 2017) was deregulated. Of note, gene ontology (GO) term analysis of the DEG showed that genes upregulated in *Usp22* KO cells were associated with inflammation and immune responses, including ISG represented by GO terms “cellular response to IFN-*β*” and “cellular response to IFN-*γ*” (Fig. 1E). Expression of individual DEG within relevant GO terms identified in Fig. 1E is shown in Fig. 1F. Gene set enrichment analysis (GSEA) revealed enhanced expression of both IFN-*α* and IFN-*γ* target genes in *Usp22* KO cells (Fig. S1B). Intersecting DEG with a comprehensive set of ISG previously found across immune cells (Mostafavi et al., 2016) identified larger numbers of ISG upregulated in *Usp22*-deficient cells (Fig. S1C). Upregulation of interferon target genes upon loss of *Usp22* was not only apparent in hematopoietic stem and progenitor cells, but also in primary pre-B cells (Fig. S1D), and pro-B cells (data not shown). In line with these findings, *Usp22* KO cells showed increased cell surface expression of MHC class II (Fig. S1E), and higher *Stat-1* (Fig. S1F), and *Ifi47* (Fig. S1G) mRNA levels, all of which are IFN-regulated. Importantly, serum concentrations of IFN-*α*, IFN-*β* and IFN-*γ* were not elevated in *Usp22* KO compared to wild type mice (Fig. 1G), and expression of the receptors for type 1 or type 2 IFNs was not increased (Fig. S1H, I). An extensive survey for pathogens found no evidence for infections in *Usp22* KO mice (Table S1). Collectively, deletion of *Usp22* in hematopoietic cells leads to enhanced monoubiquitination of H2B, and increased expression of inflammation-associated genes, including ISG. These phenotypes were observed in mice lacking signs of infections or diseases, and in the absence of elevated interferon levels.

### *Usp22* deficiency mimics interferon-related adaptive immunity

Interferons have pleiotropic effects on adaptive immunity, notably augmenting B cell responses and antibody production, and activation of T cells (reviewed in Ivashkiv and Donlin, 2013). In line with their overall IFN response-like phenotype, numbers of germinal center (GC) B cells were increased in the spleens of *Usp22* KO mice (Fig. 2A, B), and plasma cell (PC) numbers were elevated in the spleen and bone marrow (Fig. 2C-E). Consistent with this increase in PC numbers, serum concentrations of IgM, IgG1, IgG2b and IgA immunoglobulin isotypes were elevated in *Usp22* KO mice (Fig. 2F). GC B cell and PC accumulation, and enhanced antibody production were recapitulated in lymphocyte specific (*Usp22^fl/fl^* x *CD127-Cre*) (Fig. S2A-E), but not B cell specific (*Usp22^fl/fl^* x *CD19-Cre*) (Fig. S2F-I) *Usp22* KO mice, arguing for a lymphocyte-intrinsic but B cell-extrinsic effect.

**Figure 2.**
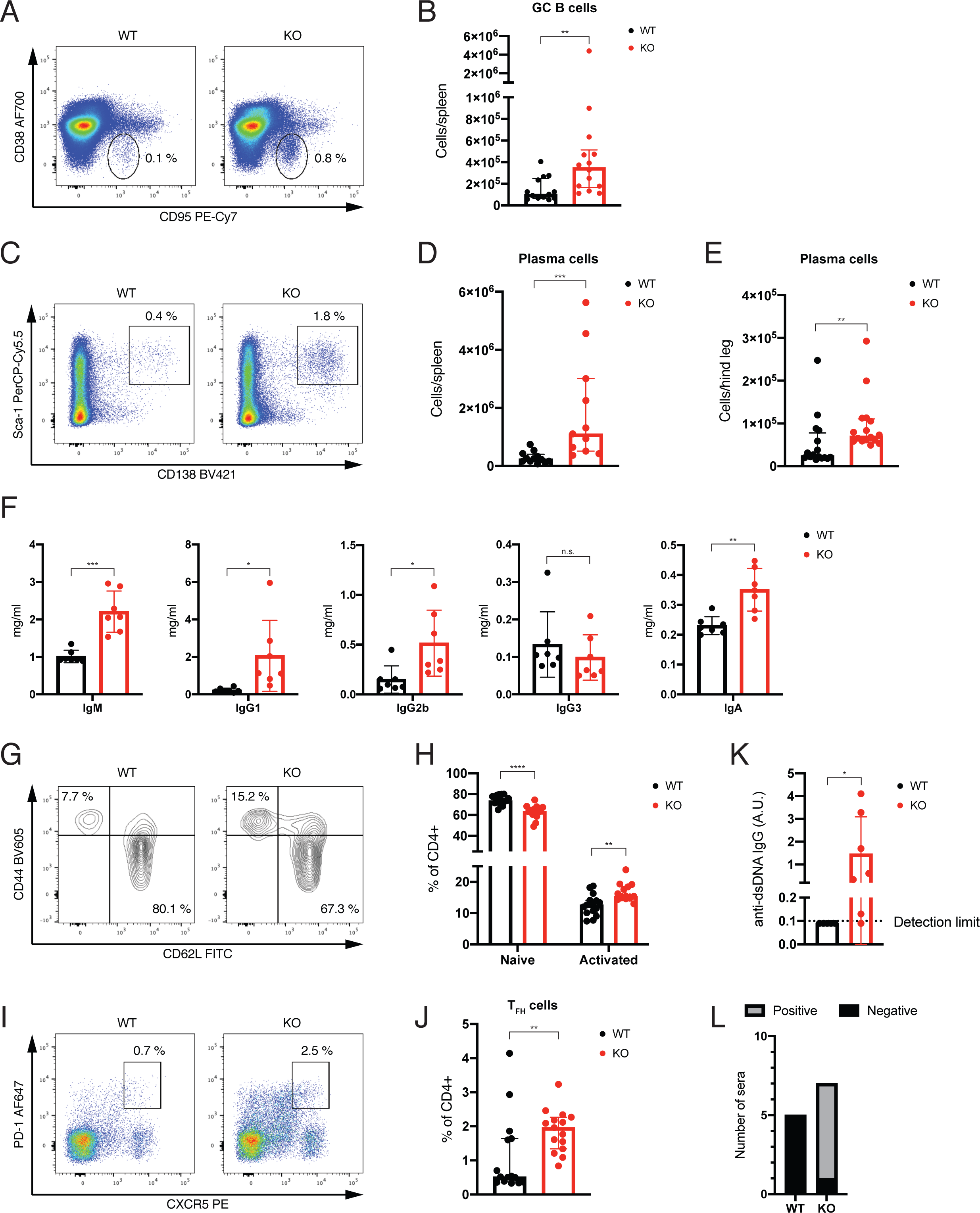
Adaptive immunity in the absence of *Usp22* resembles interferon responses. (**A**) Percentages of CD38^-^CD95^+^ germinal center (GC) B cells (pre-gated on CD19^+^B220^+^ B cells) comparing wild type and *Usp22* KO spleens. (**B**) Absolute numbers of GC B cells (n=13-14) in the spleens of wild type and *Usp22* KO mice. (**C**) Percentages of Sca-1^+^CD138^high^ plasma cells (PC) in wild type and *Usp22* KO spleens. (**D** and **E**) Absolute numbers of plasma cells in spleens (n=11-12) (**D**), and bone marrow (n=16) (**E**) of wild type and *Usp22* KO mice. (**F**) Serum concentrations of the indicated immunoglobulin isotypes in wild type and *Usp22* KO mice (n=7). (**G**) Percentages of CD62L^+^CD44^low^ naïve and CD62L^-^CD44^high^ activated T cells (pre-gated on CD3^+^CD4^+^) in wild type and *Usp22* KO spleens. (**H**) Relative proportions of naïve and activated CD4^+^ T cells (gated as in **G**) in the spleens of wild type and *Usp22* KO mice (n=15). (**I**) Percentages of PD-1^+^CXCR5^+^ T follicular helper (TFH) cells (pre-gated on CD3^+^CD4^+^) in wild type and *Usp22* KO spleens. (**J**) Percentages of TFH cells (gated as in **I**) within the splenic CD4^+^ T cell populations of wild type and *Usp22* KO mice (n=15). (**K**) Serum levels of anti-dsDNA autoantibodies detected by ELISA in wild type and *Usp22* KO mice (n=5-7) (A.U. = arbitrary units). (**L**) Fraction of sera from wild type and *Usp22* KO mice reactive (positive) and non-reactive (negative) against HEp-2 cells (n=5-7). Median ± interquartile range (**B**, **D**, **E**, **H**, **J**), mean ± s.d. (**F**, **K**). * p<0.05; ** p<0.01; *** p<0.001; **** p<0.0001; n.s., not significant by unpaired, two-tailed Mann-Whitney test (**B**, **D**, **E**, **H**, **J**), unpaired, two-tailed Student’s t-test (**F**) or unpaired, one-tailed Student’s t-test (**K**).

T cell analysis uncovered spontaneous T cell activation upon pan-hematopoietic and lymphocyte-specific deletion of *Usp22* (Fig. 2G, H; Fig. S2J, K), Importantly, *Usp22* deletion in lymphocytes resulted in enhanced differentiation of naïve T cells into CXCR5^+^PD-1^+^ T follicular helper (TFH) cells (Fig. 2I, J; Fig. S2L, M). Given the central role of TFH cells in the germinal center reaction, their accumulation matches the increased numbers of germinal center B cells, plasma cells and the enhanced antibody production. Finally, we found evidence for autoantibody production, including reactivity against double stranded DNA (Fig. 2K), and HEp-2 cells (Fig. 2L), consistent with the known link between chronic IFN responses and autoimmunity (Crow et al., 2019). Taken together, our results indicate that *Usp22* deficiency phenocopies typical IFN-stimulated adaptive immunity.

### *Usp22* deficiency mimics interferon stimulation in hematopoiesis in vivo

Interferon stimulation has profound effects on bone marrow hematopoiesis, including cell cycle activation of HSC (Essers et al., 2009; Baldridge et al., 2010), increased myeloid output (Buechler et al., 2013) and impaired B cell development (Lin et al., 1998). Given that *Usp22* KO cells showed a marked ISG expression signature, we analyzed *Usp22* KO mice for interferon phenotypes in hematopoiesis. *Usp22* KO mice had decreased absolute numbers of phenotypic long-term hematopoietic stem cells (LT-HSC), accompanied by markedly increased proliferation (Fig. S3A, B). The next hallmark of hematopoiesis in *Usp22* KO bone marrow were elevated numbers of myeloid cells (Fig. 3A), and reduced numbers of B cells (Fig. 3B). Immature granulocytes, characterized by low expression of Gr-1/Ly6G, were enriched (Fig. 3C, D; Fig. S3C, D). Consistent with inflammatory myelopoiesis, numbers of macrophages and Ly6C^high^ inflammatory monocytes, but not Ly6C^low^ patrolling monocytes, were increased in *Usp22* KO bone marrow (Fig. S3E).

**Figure 3.**
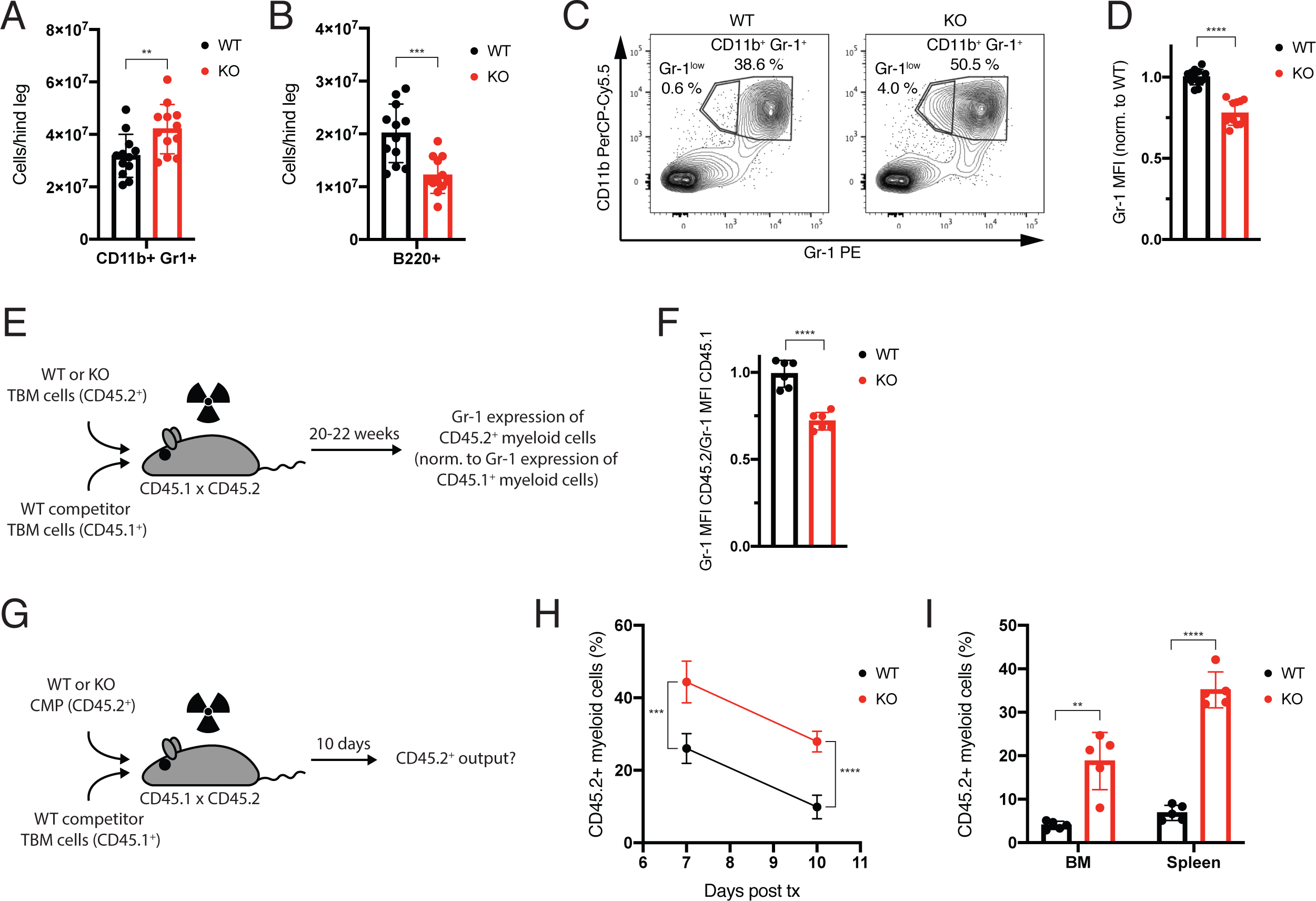
Enhanced myeloid development in *Usp22* KO mice. (**A**) Absolute numbers of CD11b^+^Gr^+^ myeloid cells in the bone marrow of wild type and *Usp22* KO mice (n=12). (**B**) Absolute numbers of B220^+^ B cells in the bone marrow of wild type and *Usp22* KO mice (n=12). (**C**) Flow cytometric analysis of wild type and *Usp22* KO BM cells stained for CD11b and Gr-1. (**D**) Gr-1 median fluorescence intensity (MFI) of CD11b^+^Gr-1^+^ BM cells from wild type and *Usp22* KO mice normalized to wild type (n=12). (**E**) Experimental setup for the generation of mixed bone marrow chimeras. (**F**) Gr-1 MFI of CD45.2^+^ test myeloid cells normalized to CD45.1^+^ competitor myeloid cells (both gated as shown in **C**) from the same recipient mouse in mixed bone marrow chimeras (n=6). (**G**) Transplantation of either wild type, or *Usp22* KO CMP (both CD45.2^+^) together with total bone marrow competitor cells (CD45.1^+^) into lethally irradiated recipients. (**H** and **I**) Relative contributions of wild type and *Usp22* KO CMP to CD11b^+^ myeloid cells in peripheral blood (**H**), and in bone marrow and spleen (**I**) (n=5). Mean ± s.d. * p<0.05; ** p<0.01; *** p<0.001; **** p<0.0001; n.s., not significant by unpaired, two-tailed Student’s t-test.

To address whether phenotypes in myelopoiesis were cell-intrinsic, *Usp22* KO bone marrow cells were transplanted competitively into lethally irradiated recipients (Fig. 3E), leading to the re-emergence of the immature granulocyte phenotype from *Usp22* KO donor bone marrow (Fig. 3F). At the progenitor level, *Usp22*-deficient common myeloid progenitors (CMP) were cell intrinsically more competitive versus total bone marrow than wild type CMP (Fig. 3G, H, I; Fig. S3F). Elevated numbers of immature granulocytes, monocytes and macrophages, and enhanced myeloid output upon adoptive transfer of CMP are all evidence for cell-autonomously enhanced myelopoiesis, a hallmark of interferon immunity, in the absence of *Usp22*.

### *Usp22* KO progenitors generate myeloid cells in lieu of B lymphocytes in vitro

To test whether myelopoiesis was favored even under conditions promoting B cell development, we cultured wild type and *Usp22* KO hematopoietic stem and progenitor cells (HSPC) on OP9 stromal cells in the presence of lymphoid growth factors, IL7 and Flt3 ligand (Nakano et al., 1994; Cho et al., 1999). Input cells were uncommitted, and lacked expression of myeloid or B cell lineage markers (Fig. 4A, upper panels). Wild type progenitors gave rise, as expected, to B220^+^CD11b^-^ B lineage cells. In marked contrast, *Usp22*-deficient progenitors yielded almost exclusively B220^-^CD11b^+^ myeloid lineage cells (Fig. 4A, lower panels, Fig. 4B). To address whether this fate bias was cell-autonomous, we set up cultures from mixed HSPC populations (50% *Usp22* wild type cells and 50% *Usp22* KO cells), and analyzed their progeny after culture. *Usp22*-deficient progenitors again made myeloid cells while the B cell fate of wild type cells was unperturbed in the presence of *Usp22*-deficient cells (Fig. 4C, D). Thus, *Usp22* KO progenitors generate myeloid cells even under B cell promoting conditions in a cell-intrinsic manner. This phenotype was not transmittable to co-cultured wild type cells. Finally, we exploited this system to analyze whether the *Usp22* KO phenotype was reversible, akin to a switch, upon re-introduction of *Usp22* into *Usp22*-deficient progenitors. Following transduction with a retroviral vector expressing *Usp22* cDNA, levels of H2Bub1 decreased (Fig. S4A), and the cells regained B cell potential (Fig. S4B, C). Hence, *Usp22* can reduce monoubiquitination of H2B within days, and the developmental bias from B to myeloid lineages is rapidly reversible.

**Figure 4.**
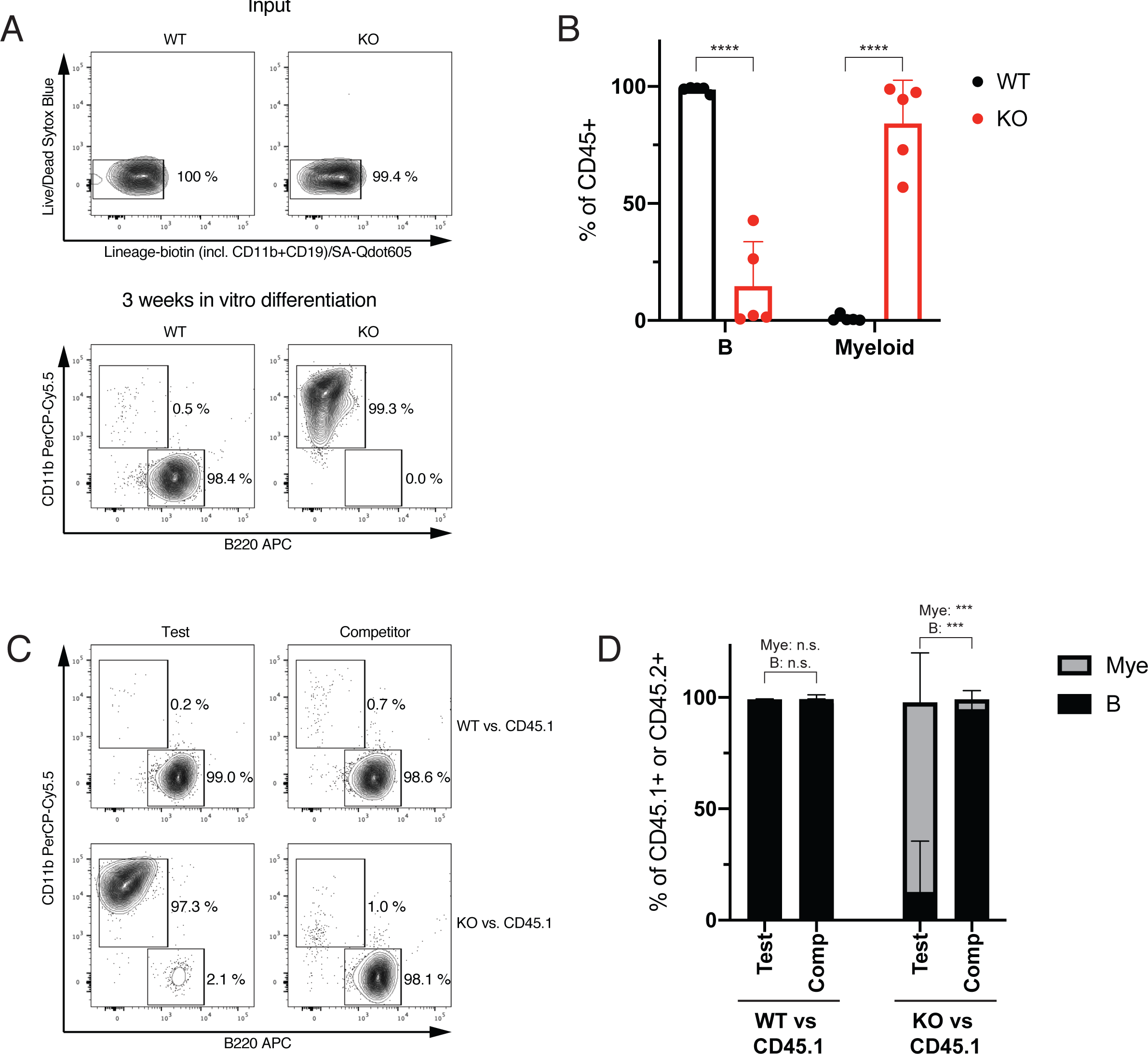
Myeloid bias of *Usp22* KO HSPC in vitro. (**A**) Absence of lineage markers CD11b (myeloid) and B220 (B lineage) on sorted wild type and *Usp22* KO HSPC before in vitro differentiation (top panels), and CD11b versus B220 expression 3 weeks after in vitro differentiation (bottom panels) (representative of 4 experiments). (**B**) Relative proportions of B220^+^ B cells and CD11b^+^ myeloid cells (gated as shown in **A**) 3 weeks after in vitro differentiation of wild type or *Usp22* KO HSPC. Each dot represents the mean of 3-5 replicates of progenitors from one donor mouse. Graph represents the results from 5 mice per genotype. (**C**) In vitro differentiation (as in **A**), mixing either wild type or *Usp22* KO HSPC (both CD45.2^+^) with wild type competitor HSPC (CD45.1^+^). After 3 weeks, cells were gated for CD45.1 versus CD45.2, and analyzed as in **A**. (**D**) Relative proportions of B220^+^ B cells and CD11b^+^ myeloid cells (gated as in **C**) in cultures derived from mixed HSPC (n=4 replicates from one mouse per genotype). Mean ± s.d. * p<0.05; ** p<0.01; *** p<0.001; **** p<0.0001; n.s., not significant by unpaired, two-tailed Student’s t-test.

### *Usp22* deficiency impairs B lymphocyte development

In keeping with the inhibition the B cell development in the bone marrow by interferons (Lin et al., 1998), total numbers of B220^+^ cells were reduced in *Usp22* KO mice (Fig. 3B). Closer inspection of stages of B cell development (Hardy et al., 1991) pinpointed the defect to fraction D (B220^+^CD43^-^IgM^-^) and onwards stages (Fig. 5A). The decrease of more mature B cells in *Usp22* KO mice is evident by the altered proportions of CD43^+^ (more immature) versus CD43^-^ (more mature) B220^+^ cells (Fig. S5A). The partial developmental block at the transition to the pre-B cell stage (fraction D) was associated with increased apoptosis of pre-B cells and following stages (Fig. 5B), again in keeping with an interferon-stimulation-like phenotype (Grawunder et al., 1993). RNA sequencing showed, in addition to enhanced ISG expression (Fig. S1D), strongly decreased expression of *Myc* and *Myc* target genes in *Usp22*-deficient pre-B cells (Fig. 5C), consistent with suppression of *Myc* mRNA expression by type I interferon-induced genes (Sarkar et al., 2006). Collectively, the stage of the developmental impairment, the increase in apoptosis, and the loss of *Myc* target gene expression all resemble the effects of *Myc* deficiency on B cell development (Vallespinós et al., 2011). The partial block in B cell development, reflected by the larger representation of CD43^+^B220^+^ cells, was recapitulated in mixed bone marrow chimeras (Fig. S5B, C), demonstrating the cell intrinsic basis of this phenotype. In the spleens, absolute numbers of B220^+^CD19^+^ B cells were reduced in *Usp22*-deficient mice (Fig. S5D), mainly resulting from a reduction in follicular I B cells (CD19^+^IgM^-^IgD^+^CD93^-^) (Fig. S5E). Collectively, *Usp22* KO mice show a cell-intrinsic incomplete block in B cell development which is, however, still permissive to generate all splenic B cell subsets.

**Figure 5.**
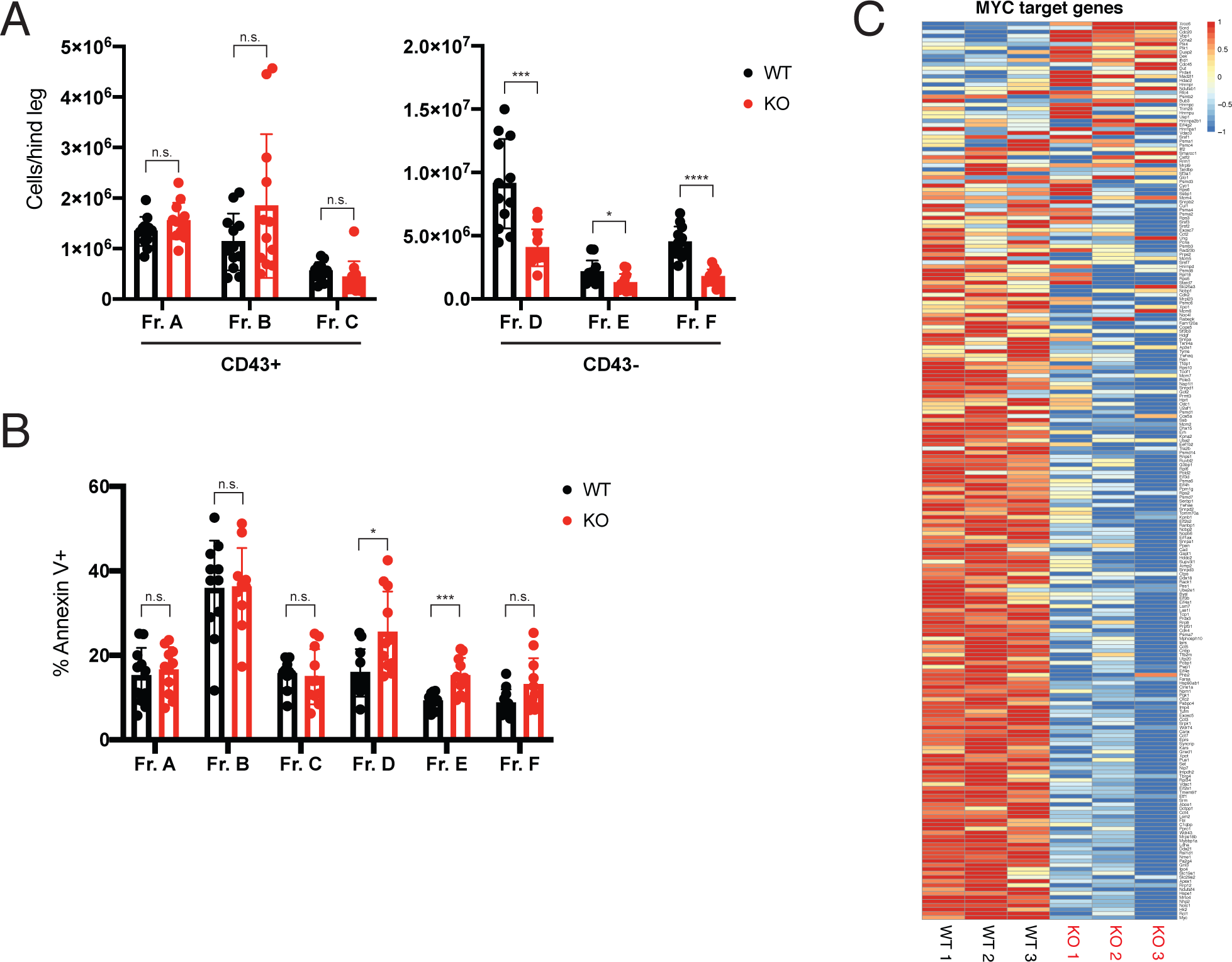
Impaired B cell development in *Usp22* KO mice. (**A**) Absolute numbers of the indicated B cell progenitor populations in the bone marrow of wild type and *Usp22* KO mice (n=12). (**B**) Percentage of Annexin V^+^ apoptotic cells within the indicated B cell progenitor populations of wild type and *Usp22* KO mice (n=12). (**C**) Heatmap showing the expression of *Myc* target genes in primary pre-B cells (Fraction C) from wild type and *Usp22* KO mice. Mean ± s.d. * p<0.05; ** p<0.01; *** p<0.001; **** p<0.0001; n.s., not significant by unpaired, two-tailed Student’s t-test.

### *Usp22* deficiency increases H2B monoubiquitination of nucleosomes at ISG loci

In order to intersect gene expression changes caused by *Usp22* deficiency with alterations in locus-specific H2Bub1 levels and chromatin accessibility, we complemented the RNA sequencing data with chromatin immunoprecipitation (ChIP) sequencing and assay for transposase accessible chromatin (ATAC) sequencing. We used ChIP sequencing to identifiy H2Bub1-bound loci in wild type and *Usp22* KO cells. Genome-wide data revealed the typical pattern of gene body H2B monoubiquitination (Shema et al., 2008), with H2Bub1 levels peaking at the beginning of the transcribed region and then decreasing towards the end of the gene body (Fig. S6A). H2B monoubiquitination was absent in the gene bodies of non-expressed genes, and high in the transcribed regions of strongly expressed genes (Fig. S6B), consistent with RNA polymerase II-dependent recruitment of the RNF20/RNF40 complex (Zhang and Yu, 2011). Both of these principle attributes of H2Bub1 were unaffected by *Usp22* deficiency (Fig. S6A, B). However, *Usp22* deficiency strongly increased H2B monoubiquitination of nucleosomes at transcribed loci (Fig. S6B, bottom panels). Consistent with the interferon-like phenotype observed in *Usp22*-deficient mice, gene body H2B monoubiquitination was also enhanced at genes normally regulated by IFN in *Usp22* KO cells (Fig. 6A, Fig. S6C).

**Fig. 6.**
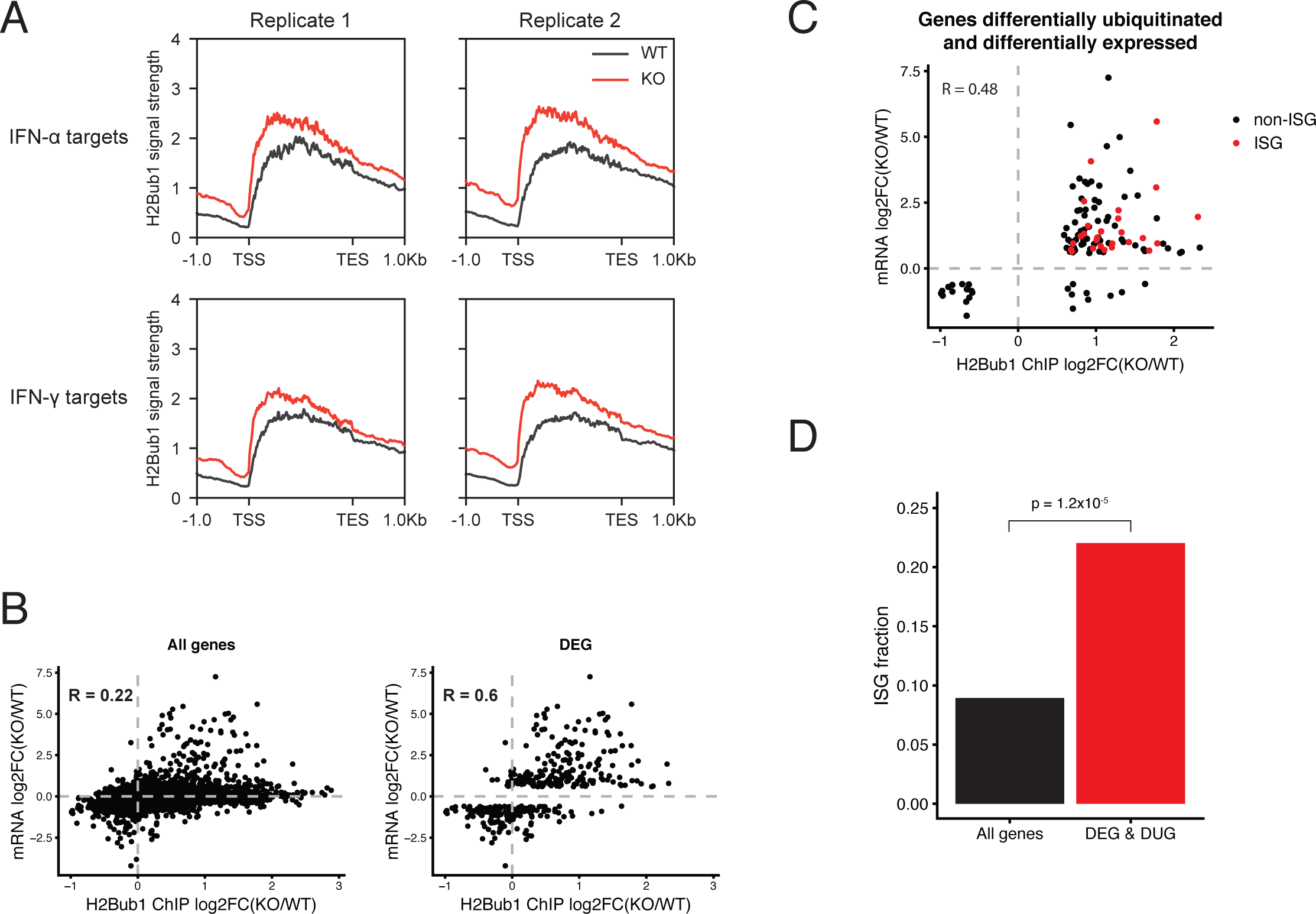
Increased H2Bub1 levels at ISG loci in *Usp22* KO cells. (**A**) Average H2Bub1 ChIP sequencing signal across gene bodies of IFN-*α* target genes (top panels) and IFN-*γ* target genes (bottom panels) in wild type and *Usp22* KO HSPC. (**B**) Correlation between change in gene body monoubiquitination and change in mRNA expression comparing *Usp22* KO against wild type HSPC for individual genes (dots), shown for all genes (left panel), or selectively for differentially expressed genes (right panel). (**C**) Correlation between change in gene body monoubiquitination and change in mRNA expression upon loss of *Usp22* in HSPC for genes differentially expressed and ubiquitinated between wild type and *Usp22* KO HPSC. ISG are highlighted in red. (**D**) Fraction of ISG (Mostafavi et al., 2016) within all genes detected in RNA sequencing and H2Bub1 ChIP sequencing (black bar), and the set of genes differentially expressed and ubiquitinated in *Usp22* KO HSPC (red bar). p-value was determined using Fisher’s exact test. TSS = Transcription start site; TES = Transcription end site.

In search for a link between altered gene expression and enhanced H2B monoubiquitination in *Usp22*-deficient cells, we integrated RNA sequencing and H2Bub1 ChIP sequencing data. Looking at all genes, the correlation between change in expression and change in gene body monoubiquitination upon loss of *Usp22* was poor (Fig. 6B, left panel). In contrast, there was good correlation between change in expression and change in gene body monoubiquitination for differentially expressed genes (Fig. 6B, right panel), arguing for a gene set-specific role of *Usp22* in the regulation of gene expression. Of note, genes downregulated in *Usp22* KO cells did not show a consistent change in gene body H2B monoubiqutination (Fig. 6B, right panel, lower quadrants), suggesting that changes in mRNA expression do not automatically result in alterations of H2Bub1 levels. Based on ATAC sequencing, genes differentially expressed between wild type and *Usp22* KO cells showed on average no changes in promoter accessibility (Fig. S6D), and for individual genes there was no correlation between change in expression and change in promoter accessibility upon loss of *Usp22* (Fig. S6E). In order to identify genes directly regulated by *Ups22* through H2B monoubiquitination, we searched for genes for which both mRNA expression and gene body H2Bub1 levels changed (either up or down). In total, we identified 118 such genes, of which 102 showed increased H2B monoubiquitination (Fig. 6C, right quadrants), and 93 exhibited both enhanced H2B monoubiquitination and increased mRNA expression in *Usp22* KO cells (Fig. 6C, upper right quadrant). When filtering differentially expressed and ubiquitinated genes for ISG (26 genes), all of these showed both enhanced expression and increased H2B monoubiquitination (Fig. 6C, red dots). Among all genes, for which we obtained information both on gene expression and gene body H2Bub1 levels, each regardless of change, approximately 8% were ISG (Mostafavi et al., 2016), roughly representing ISG abundance in the genome (Fig. 6D, all genes). By comparison, genes that were both differentially expressed and ubiquitinated in *Usp22*-deficient cells were enriched almost 3-fold for ISG (Fig. 6D, DEG & DUG), implying preferential regulation of ISG by *Ups22* and H2B monoubiquitination levels. Of note, both the group of all differentially expressed and differentially ubiquitinated genes (mostly upregulated and hyperubiquitinated), and the group of differentially expressed and differentially ubiquitinated ISG (exclusively upregulated and hyperubiquitinated) showed no change in promoter accessibility (Fig. S6F). Taken together, these data demonstrate that USP22 acts as a deubiquitinase for monoubiquitinated H2B at ISG loci. Higher H2Bub1 levels were observed at more than half of the upregulated ISG loci (26 of 48) (Fig. 6C, Fig. S1C). Enhanced ISG expression in *Usp22* KO cells was independent of increased promoter accessibility.

## Discussion

Combining immunophenotyping with transcriptome and epigenome analysis of three different stage-and lineage-specific *Usp22* knock-out mouse lines, we identified *Usp22* as a molecular switch that coordinates diverse intracellular and systemic interferon responses in vivo. Interferons are secreted by most cell types upon viral or bacterial infections, and play pivotal roles in host defense. Engagement of the type I IFN receptor results in phosphorylation of STAT proteins and formation of STAT homo- or heterodimers that translocate into the nucleus. Depending on their composition and association with additional factors, such as IRF9, STAT dimers can bind to the promoters of different classes of ISG and activate their transcription (Ivashkiv and Donlin, 2013; Schneider et al., 2014). While processing and relay of intracellular IFN signaling is multifaceted, it is unknown whether an epigenetic state exists that elicits downstream ISG expression, ultimately driving IFN-associated innate and adaptive immune responses.

While previous data showed in a cell line that H2Bub1 is required for ISG expression (Fonseca et al., 2012), a key question was whether higher levels of H2B monoubiquitination are sufficient to drive ISG expression. If so, the enzymes modulating the degree of H2B monoubiquitination might allow to regulate intracellular and systemic IFN responses, with the E3 ubiquitin ligase RNF20 catalyzing, and the H2B deubiquitinase USP22 reversing H2B monoubiquitination. We observed stronger H2B monoubiquitination in *Usp22*-deficient immune cells in mice which is consistent with a perturbed balance towards higher H2B monoubiquitination in the absences of this deubiquitinase (Li et al., 2018; Melo-Cardenas et al., 2018; Cortez et al., 2020). *Usp22* deficiency resulted not only in increased H2Bub1 levels, but also enhanced ISG expression. Consistent with upregulated ISG expression, pan-hematopoietic *Usp22* KO mice exhibited hallmarks of the systemic IFN response, including increased HSC proliferation, enhanced myelopoiesis, impaired B cell development, spontaneous T cell activation, increased numbers of germinal center B cells and plasma cells, elevated immunoglobulin serum concentrations and the presence of autoantibodies. Considering the broad range of IFN-driven processes affecting multiple arms of immunity, it seems remarkable that most, if not all, can be elicited by loss of *Usp22*. It remains to be determined whether these broad effects are mediated by cell type-specific or universal transcriptional changes driven by the loss of *Usp22*. In the context of a virus infection, *Usp22* regulates IRF3 localization and activity, a transcription factor driving the expression of IFN-*β* (Cai et al., 2020). However, our results indicate that increased ISG expression, and IFN-related immunity upon *Usp22* deletion develop in the absence of detectable signs of bacterial or viral infections and elevated serum IFN levels. Altered hematopoiesis (enhanced myelopoiesis, myeloid bias, and impaired B cell development) occurred in a cell-autonomous manner in mixed bone marrow chimeras, and in co-culture experiments in vitro, arguing against the involvement of a secreted factor that would transfer the phenotype from *Usp22* KO to wild type cells. In contrast to the cell-autonomous effects described before, the phenotypes affecting the adaptive immune system (T cell activation, GC B cell and PC numbers, immunoglobulin levels) were dependent on lymphocyte-but not B cell-specific *Usp22* deletion, consistent with a recent report demonstrating that *Usp22* stabilizes FOXP3, and thereby restricts anti-tumor T cell responses as well as autoimmune reactions in a regulatory T cell-intrinsic manner (Cortez et al., 2020). Our data broaden these findings as they demonstrate that lymphocyte-specific *Usp22* deficiency leads to enhanced immunoglobulin production, most likely due to increased numbers of T follicular helper cells. Given that these processes ultimately result in the production of autoantibodies, further extending the function of *Usp22* as a gatekeeper of autoimmunity, it remains to be determined how the broadly ’pre-activated’ interferon response by loss of *Ups22* affects immunity beyond anti-tumor responses (Cortez et al., 2020).

Mechanistically, in *Usp22*-deficient cells, about half of the ISG loci with enhanced expression showed increases nucleosomal H2B monoubiquitination. ISG with stronger expression upon loss of *Usp22* did not exhibit higher promoter accessibility, implying that expression changes driven by *Usp22* deficiency and increased H2Bub1 levels are likely caused by events subsequent to transcription factor binding and RNA polymerase II recruitment. H2Bub1 has been implicated in the positive regulation of transcriptional elongation (Pavri et al., 2006), which might account for the observed transcriptome changes in *Usp22*-deficient cells. It thus appears that downstream from transcription factor binding H2B monoubiquitination serves as an amplifier of expression of large numbers of ISG. According to this view, *Usp22* is a negative regulator of ISG expression in vivo. These novel insights into the broad immunological functions of *Usp22* might be relevant for the treatment of inflammatory diseases, autoimmunity and leukemias as well as the improvement of vaccination strategies.

## Materials and methods

### Mice

*Usp22* flox mice (B6-Usp22^tm1c(KOMP)Wtsi^) were crossed to *Vav1-Cre* (B6.Cg-*Commd10^Tg(Vav1-icre)A2Kio^*/J) (de Boer et al., 2003), *CD127-Cre* (B6;129-Il7r^tm1.1(icre)Hrr^) (Schlenner et al., 2010) or *CD19-Cre* mice (B6.129P2(C)-*Cd19^tm1(cre)Cgn^*/J) (Rickert et al., 1997) to obtain *Usp22* KO, *Usp22* lyKO or *Usp22* bKO mice. B6.SJL-Ptprc^a^ Ptprc^b^ mice, used as recipients in competitive transplantation experiments, were derived by inter-crossing C57Bl/6J mice (carrying the Ptprc^b^/CD45.2 allele) with B6-Ptprc^a^ mice (carrying the Ptprc^a^/CD45.1 allele). Mice were kept in individually ventilated cages under specific pathogen-free conditions in the animal facility of the German Cancer Research Center (DKFZ, Heidelberg). For most experiments, Cre-positive mice carrying two *Usp22* wild type alleles were used as wild type controls. Both male and female mice were used. No randomization and no blinding were done. All animal experiments were performed according to institutional and governmental regulations, and were approved by the Regierungspräsidium Karlsruhe, Germany.

### Bone marrow and CMP transplantation

Recipient mice were lethally irradiated with a split dose of 2 x 5.5 Gy using a Cs137 Gammacell 40 irradiator with a time interval of three to four hours between the two irradiation cycles. Donor cells (in PBS) were injected into the tail veins. For competitive bone marrow transplantations, 1x10^6^ test bone marrow cells (derived from *Usp22* KO mice or wild type littermates, both expressing CD45.2) were transplanted together with 1x10^6^ competitor bone marrow cells (derived from B6-Ptprc^a^ mice) into the same recipient. For CMP transplantations, 1x10^4^ test CMP (derived from *Usp22* KO mice or wild type littermates, both expressing CD45.2) were transplanted together with 1x10^5^ competitor bone marrow cells (derived from B6-Ptprc^a^ mice) into the same recipient. Mice were kept on Neomycin containing drinking water for two weeks after transplantation.

### Blood and serum collection

Blood samples were collected by puncture of the submandibular vein, or by post mortem cardiac puncture. For flow cytometry, blood samples were collected into EDTA-coated tubes, and subjected to red blood cell lysis using RBC lysis buffer (BioLegend) before further processing as described below. For serum preparation, blood was collected into serum tubes, incubated for 30 minutes at room temperature to allow for coagulation, and centrifuged at 15,000g for 90 seconds. Serum samples were frozen in aliquots at -80°C.

### Preparation of single cell suspensions from hematopoietic organs

Bones (tibiae, femurs, pelvis and spine) freed of other tissues were crushed in FACS buffer (PBS + 5 % FBS) using mortar and pestle, and filtered through 40 μm cell strainers into 50 ml Falcon tubes. Spleens were mashed with FACS buffer through pre-wetted 40 μm cell strainers into 50 ml Falcon tubes using syringe plungers. Cell numbers were determined using a Cellometer Auto 2000 Cell Viability Counter (Nexcelom).

### Flow cytometry and fluorescence activated cell sorting (FACS)

Single cell suspensions, or lysed blood samples were blocked for 15 minutes in 10 % normal rat serum (diluted in FACS buffer) and stained for 20 minutes in titrated dilutions of fluorescent dye-labeled antibodies (in FACS buffer) at 4°C. Live/dead cells were discriminated by Sytox Blue (Thermo Fisher Scientific). For the sorting of HSPC, CMP and B cell progenitors, bone marrow cells were subjected to lineage depletion prior to further antibody staining. Cells were labeled with biotinylated antibodies directed against lineage markers (CD4, CD8, CD11b, CD19 [omitted in the enrichment of B cell progenitors], Gr-1 and Ter119), and were depleted using anti-biotin microbeads (Miltenyi) on LS MACS columns (Miltenyi). Samples were analyzed on a LSRFortessa flow cytometer (BD Biosciences), or sorted on a FACSAria III cell sorter (BD Biosciences). Antibodies and second step reagents were: BP-1-PE (clone BP-1), CD4-biotin (GK1.5), CD4-FITC and CD4-PE (H129.19), CD8a-FITC and CD8a-PE (53-6.7), CD19-biotin (1D3), CD43-APC and CD43-biotin (S7), CD45-FITC (30-F11), CD45.2-FITC (104), CD45R-APC and CD45R-FITC (RA3-6B2), CD93-BV421 (AA4.1), CD95-PE-Cy7 (Jo2), CD244.2-FITC (2B4), Gr-1-APC and Gr-1-PE (RB6-8C5), IgM-PE (R6-60.2), Kit-APC (2B8), Ly6G-PerCP-Cy5.5 (1A8), Ter119-PE (TER-119) and Streptavidin-APC-Cy7 from BD Biosciences, CD3-BV421 (17A2), CD4-BV421 (GK1.5), CD8a-BV421 (53-6.7), CD11b-BV421 (M1/70), CD19-BV421 and CD19-BV605 (6D5), CD21/35-PerCP-Cy5.5 (7E9), CD44-BV605 (IM7), CD45.2-AlexaFluor488 (104), CD48-AlexaFluor700 (HM48-1), CD115-BV605 (AFS98), CD138-BV421 (281-2), CD150-PE and CD150-PE-Cy7 (TC15-12F12.2), CD229-PE (Ly9ab3), CXCR5-PE (L138D7), Gr-1-BV421 (RB6-8C5), IFNAR-1-PE (MAR1-5A3), IgD-APC and IgD-APC-Cy7 (11-26c.2a), Kit-BV711 (2B8), Ly6C-PE (HK1.4), MHCI-PE (28-8-6), PD-1-AlexaFluor647 (29F.1A12) and Ter119-BV421 (TER-119) from BioLegend, BP-1-biotin (6C3), CD3-APC-eFluor780 (17A2), CD8a-biotin and CD8a-PE-Cy7 (53-6.7), CD11b-biotin (M1/70.15), CD11b-PE, CD11b-PE-Cy7, CD11b-PerCP-Cy5.5 (M1/70), CD16-PE-Cy7 (93), CD23-PE-Cy7 (B3B4), CD24-eFluor780 (M1/69), CD34-eFluor660 (RAM34), CD38-AlexaFluor700 (90), CD45-PE-Cy7 (30-F11), CD45.1-PE-Cy7 (A20), CD45.2-APC (104), CD45R-PerCP-Cy5.5 (RA3-6B2), CD48-APC and CD48-FITC (HM48-1), CD62L-FITC (MEL-14), F4/80-APC and F4/80-PE (BM8), Gr-1-biotin (RB6-8C5), IFNGR-1-PE (2E2), IgM-FITC (eB121-15F9), IgM-PE-Cy7 (II/41), Kit-APC-eFluor780 and Kit-PE (2B8), MHCII-APC and MHCII-APC-eFluor780 (M5/114.15.2), Sca-1-PE-Cy7 and Sca-1-PerCP-Cy5.5 (D7), Ter119-biotin and Ter119-FITC (TER-119), Streptavidin-APC and Streptavidin-Qdot605 from Thermo Fisher Scientific.

### EdU labeling

Mice received 100 μg EdU by intraperitoneal injection and were sacrificed two hours later. Bone marrow cells were isolated, depleted and stained as described above with the modification that Zombie Red dye (BioLegend) was used for live/dead cell discrimination. Afterwards, cells were subjected to Click chemistry based EdU detection using the Click-iT Plus EdU Alexa Fluor 488 Flow Cytometry Kit (Thermo Fisher Scientific) according to the manufacturer’s instructions. Samples were labeled with FxCycle Violet stain to assess DNA content, and data were acquired on a LSRFortessa flow cytometer (BD Biosciences).

### Annexin V labeling

Bone marrow cells were isolated and stained as described above. Annexin V labeling was performed using the FITC Annexin V Apoptosis Detection Kit (BD Biosciences) according to the manufacturer’s instructions with minor modifications. 10^6^ bone marrow cells were labeled using 3 μl of FITC Annexin V. Sytox Blue was used for live/dead cell discrimination. Data were acquired on a LSRFortessa flow cytometer (BD Biosciences).

### Measurement of interferon serum levels

Serum levels of IFN-*α*, IFN-*β* and IFN-*γ* were determined using the LEGENDplex mouse type 1/2 interferon panel (BioLegend) according to the manufacturer’s instructions. Samples were analyzed on a LSRFortessa flow cytometer (BD Biosciences).

### Measurement of immunoglobulin serum levels

IgM, IgG1, IgG2b, IgG3 and IgA serum levels were determined using the LEGENDplex mouse immunoglobulin isotyping panel (BioLegend) according to the manufacturer’s instructions. Samples were analyzed on a LSRFortessa flow cytometer (BD Biosciences).

### Anti-dsDNA autoantibody ELISA

Levels of anti-dsDNA IgG were measured by ELISA using MaxiSorp plates (Nunc) coated with dsDNA from calf thymus (20 µg/ml; Sigma-Aldrich). Sera were added in 1:2 serial dilutions starting at 1:100. A serum pool obtained from 9-month-old diseased NZB/W mice served as internal standard. The starting dilution of 1:200 of the NZB/W serum was arbitrarily assigned to 100 relative units. Goat-anti-mouse IgG, Fc-specific, coupled to horseradish peroxidase was used for detection (Jackson ImmunoResearch).

### Anti-nuclear antibody (ANA) detection

ANA detection was done by indirect immunofluorescence on HEp-2 cells. Briefly, slides with fixed HEp-2 cells (EUROIMMUN, Luebeck, Germany) were incubated with 1:50 diluted sera and bound IgG was detected with Cy3-labeled goat anti-mouse IgG, Fc-specific (Jackson ImmunoResearch).

### RNA extraction

RNA was extracted from sorted cells using the Arcturus PicoPure RNA Isolation Kit (Thermo Fisher Scientific) according to the manufacturer’s instructions including the on-column DNase digestion step.

### qRT-PCR

cDNA was synthesized from RNA using the SuperScript Vilo cDNA synthesis kit (Thermo Fisher Scientific) according to the manufacturer’s instructions. Oligo-dT primers were additionally added to the reaction mix. cDNA was diluted and used for qRT-PCR using SYBR Green PCR Master Mix (Thermo Fisher Scientific) on a Viia7 Real-Time PCR System (Applied Biosystems). Fold change expression values were determined with the comparative delta Ct method using *Sdha* as reference gene. Primer sequences are given in Table S2. Forward and reverse primers used for detection of *Usp22* mRNA were located in exon 7 and exon 8, and hence downstream of the loxP site-flanked exon 2.

### RNA sequencing

Equal amounts of total RNA were converted into double-stranded cDNA using the SMARTer Ultra Low Input RNA for Illumina Sequencing – HV kit according to the manufacturer’s instructions. Sequencing libraries were prepared with the NEBNext ChIP-Seq Library Prep Master Mix Set for Illumina according to the manufacturer’s instructions with the following modifications. 10 μl of adapter-ligated double-stranded cDNA were amplified using NEBNext Multiplex Oligos for Illumina (25 µM primers), NEBNext High-Fidelity 2x PCR Master Mix and 15 cycles of PCR. Libraries were validated using an Agilent 2200 TapeStation and a Qubit fluorometer. 50 bp single-read sequencing of pooled libraries was performed on an Illumina HiSeq 2000 v4.

### OMNI-ATAC sequencing

OMNI-ATAC sequencing was performed as described in (Corces et al., 2017) with minor modifications. Briefly, 50,000 sorted HSPC were lyzed and nuclei subjected to Tn5 mediated transposition for 30 minutes at 37°C. DNA was purified using Ampure XP beads and used for indexing PCR. Sequences of the used indexing primers are given in Table S3. The number of PCR cycles was determined by qRT-PCR as described in (Buenrostro et al., 2015). Amplified libraries were purified by two-sided Ampure XP bead purification and validated on a Bioanalyzer (Agilent). 50 bp paired-end sequencing of pooled libraries was performed on an Illumina HiSeq2500.

### ChIPmentation for H2Bub1

H2Bub1 ChIPmentation of HSPC was performed according to (Schmidl et al., 2015) with minor modifications. Therefore, sorted HSPC were subjected to cross-linking in 1 % formaldehyde (in PBS) for 10 minutes at room temperature, and cross-linking was stopped by adding glycine. Pellets of cross-linked cells were resuspended in shearing buffer (10 mM Tris-HCl pH 8, 0.1% SDS, 1 mM EDTA) supplemented with Complete Protease Inhibitor Cocktail (Roche), and chromatin was sheared to an average fragment size of 200-250 bp. DNA concentrations were assessed using a Qubit fluorometer, equal amounts of sheared chromatin were diluted in ChIPmentation dilution buffer (20 mM HEPES pH 8.0, 150 mM NaCl, 0.1 % SDS, 1 % Triton-X, 1 mM EDTA, 0.5 mM EGTA), and incubated with anti-H2Bub1 (clone D11, Cell Signaling Technologies, order no. 5546S) or rabbit isotype control (BioLegend, order no. 910801) antibody over night at 4°C. On the next day, pre-blocked magnetic Protein A beads were added, and samples incubated for two hours at 4°C. Bead-bound chromatin fragments were separated on a magnetic tube rack and washed before they were subjected to tagmentation. Chromatin fragments bound to magnetic beads were resuspended in 30 μl Tagmentation mix (14 μl H2O, 15 μl Tagment DNA Buffer [Illumina, order no. 15027866], 1 μl Tagment DNA Enzyme [Illumina, order no. 15027865]) and incubated for 10 minutes at 37°C. Bead-bound chromatin was washed and decross-linked in elution buffer containing Proteinase K for eight hours at 56°C. Decross-linked chromatin was purified using Ampure XP beads, subjected to indexing PCR as described in the section “OMNI-ATAC sequencing” and purified again with Ampure XP beads. Purified libraries were validated on a Bioanalyzer instrument. 50 bp single-end sequencing of pooled libraries was performed on an Illumina HiSeq2500.

### Single-cell *Usp22* genotyping

Single cells were sorted directly into PCR tubes containing 25 μl PCR lysis buffer (1x Buffer 1 from the Expand Long Template PCR System [Roche], 0.5 mg/ml Proteinase K) and lysed for 1h at 55°C. Proteinase K was heat-inactivated for 10 min at 95°C before 25 μl 2x PCR mix were added for the first amplification. 2 μl of the first PCR reaction were used as template for the second round of nested PCR amplification. Primer sequences are given in the Key Resources Table.

### Western Blot

Cell pellets were resuspended in RIPA buffer supplemented with Complete Protease Inhibitor Cocktail (Roche), incubated for 30 minutes on ice and sonicated using a Bioruptor Plus sonicator. Samples were cleared of debris by centrifugation. Cell lysates were supplemented with Laemmli Buffer, heat-denatured, and separated by SDS-PAGE. Equal cell numbers were loaded. Proteins were transferred to PVDF membranes. Membranes were blocked in 5 % milk (in TBS-Tween) and probed with primary antibodies overnight. Primary antibodies were detected using HRP-conjugated secondary antibodies in combination with ECL substrate solution. Due to differences in their signal strengths, different exposure times were chosen for the detection of H2B and H2Bub1. Antibodies used were: anti-H2B (clone mAbcam 52484, Abcam, order no. 52484), anti-H2Bub1 (clone D11, Cell Signaling Technologies, order no. 5546S), HRP-conjugated goat-anti-mouse IgG (Thermo Fisher Scientific, order no. 32430), HRP-conjugated goat-anti-rabbit IgG (Thermo Fisher Scientific, order no. 32460). Results shown in Fig. 1D are representative of n=8 mice/genotype analyzing whole tissue lysates from various hematopoietic organs (bone marrow, spleen and thymus) or lysates of purified hematopoietic cell types.

### OP9 assay

Sorted HSPC (500 cells/well on 24-well plates) were plated onto irradiated OP9 feeder cells in B cell differentiation medium (MEM*α* + 20 % FBS + Penicillin/Streptomycin + 10 ng/ml rmFlt3L + 5 ng/ml rmIL-7) and cultured for 1.5 to 3 weeks. At the end of the differentiation period, cells were collected by pipetting and stained for flow cytometry as described above.

### Gamma-retroviral (GRV) transduction of HSPC

The GRV backbone pMIG was a gift from William Hahn (Addgene plasmid # 9044). GRV particles were produced by transient transfection of Phoenix GP cells and concentrated using Retro-X concentrator (Takara). Sorted HSPC were pre-expanded for two days, spin-infected (multiplicity of infection = 15) in the presence of polybrene and subjected to OP9 in vitro differentiation assays.

### RNA sequencing data processing and analysis

The RNA sequencing reads were first subjected to adapter trimming and low-quality read filtering using *flexbar* (version 2.5) (Dodt et al., 2012) with the following parameters: -u 10 -m 32 -ae RIGHT -at 2 -ao 2. Reads that were mapped to the reference sequences of rRNA, tRNA, snRNA, snoRNA, and miscRNA (available from Ensembl and RepeatMasker annotation) using *Bowtie* 2 (version 2.2.9) (Langmead and Salzberg, 2012) with default parameters (in --end-to-end & --sensitive mode) were excluded. The remaining reads were then mapped to the mouse reference genome (mm10) using *Tophat2* (version 2.1.1) (Kim et al., 2013) with the parameters -N 2 --read-gap-length 2 --read-edit-dist 3 --min-anchor 6 --library-type fr-unstranded --segment-mismatches 2 --segment-length 25, or *STAR* (version 2.7.3a) (Dobin et al., 2012) with key parameters --outFilterMismatchNmax 8 --outFilterMismatchNoverLmax 0.04 --alignIntronMin 20 --alignIntronMax 100000 -- outFilterType BySJout --outFilterIntronMotifs RemoveNoncanonicalUnannotated. Reads that were mapped to multiple genomic sites were discarded in the following analysis. *HTSeq-count* (version 0.9.1) (Anders et al., 2015) was used to count reads mapped to annotated genes, with parameters -f bam -r pos -s no -a 10.

Differentially expressed gene analysis was performed with the R package *DESeq2* (version 1.16.0) (Love et al., 2014). In brief, size factor estimation was first conducted to normalize the data across samples, followed by dispersion estimation to account for the negative binomial distributed count data in RNA sequencing. Finally, gene expression fold changes were calculated and the significance of the gene expression difference was estimated using the Wald test. To control the false discovery rate in multiple testing, the raw p-values were adjusted using the Benjamini–Hochberg (BH) procedure. Genes with adjusted p-values less than 0.05 and fold changes over 1.5 folds were considered as differentially expressed.

### ChIP sequencing data processing and analysis

The ChIP sequencing reads were first subjected to adapter trimming and low-quality read filtering using *flexbar* (version 2.5) (Dodt et al., 2012) with the following parameters: -u 5 -m 26 -ae RIGHT -at 2 -ao 1. The trimmed reads were mapped to the mouse reference genome (mm10) using *Bowtie* 2 (version 2.2.9) (Langmead and Salzberg, 2012) with default parameters. Reads that mapped to mitochondrial DNA or with low mapping quality (< 30) were excluded for downstream analysis. Duplicate reads due to PCR amplification of single DNA fragments during library preparation were identified using *Picard* (version 2.17.3; available at http://broadinstitute.github.io/picard) and thus removed from the downstream analysis.

To obtain the pure signal of monoubiquitinated H2B, H2Bub1 ChIP sequencing data were compared to the input data (libraries prepared from DNA of cells from replicate set 1) using the *bdgcmp* command of *macs2* (version 2.1.1) (Zhang et al., 2008b) for signal fold enrichment. The resulting signal was then used for drawing signal profiles and heatmaps. To identify differentially ubiquitinated genes, we counted the deduplicated reads overlapping with gene bodies (introns included) of individual genes. *DESeq2* (version 1.16.0) (Love et al., 2014) was then used for statistical comparison of the read counts obtained from wild type and *Usp22*-deficient HSPC, with a procedure similar to the analysis of the RNA sequencing data. Genes with adjusted p-values less than 0.05 and fold changes over 1.5 folds were considered as differentially ubiquitinated.

### ATAC sequencing data processing and analysis

The processing steps of ATAC sequencing data for read trimming, mapping and deduplication were the same as those for the ChIP sequencing data. Genome-wide chromatin accessibility was quantified using *bamCoverage* from the *deeptools* python toolkit suit (version 3.4.3) (Ramírez et al., 2016) based on the deduplicated reads. To compare data from different libraries, the accessibility signal was normalized using library scale factors. Thus, the normalized accessibility signal was used for drawing signal profiles and heatmaps. To identify genes with differential accessibility in their promoters, we counted the deduplicated reads overlapping with gene promoters (defined as a 2000-bp window centered at a gene’s transcription start site). *DESeq2* (version 1.16.0) (Love et al., 2014) was then used for statistical comparison of the read counts obtained from wild type and *Usp22*-deficient HSPC, with a similar procedure as for analyzing the RNA sequencing data. Genes with adjusted p-values less than 0.05 and fold changes over 1.5 folds were considered as the ones with differential promoter accessibility.

### GO/hallmark signature enrichment analysis

The gene symbols were mapped to GO terms/hallmark gene sets using either R packages *GO.db*, *AnnotationDbi*, and *org.Mm.e.g.db*, or based on the *MSigDB* collections (Liberzon et al., 2015). Gene sets with at least 10 genes in background (i.e. all expressed genes) were tested for enrichment in DEG using the GOseq method provided in the R package *goseq* (Young et al., 2010). The raw p-values were then adjusted using the Benjamini– Hochberg (BH) procedure. Gene set enrichment analysis (GSEA) (Subramanian et al., 2005) was performed using the GSEA JAVA interface (version 4.0.3) available at https://www.gsea-msigdb.org/gsea/.

### Statistical analysis

Graphs were prepared using GraphPad Prism or R. Statistical details, including definition of center and precision measures, number of replicates (n) as well as statistical tests used can be found in the figure legends. For normally distributed data, mean ± standard deviation (s.d.) is shown and statistical significance was tested using Student’s t-test. For data that did not follow a normal distribution, median ± interquartile range is shown and significance was tested using Mann-Whitney test.

## Acknowledgments

We thank Sandrine Sander and Klaus Rajewsky for generously providing *CD19*-Cre mice. We thank the Genomics and Proteomics Core Facility for providing Illumina sequencing services, Annette Kopp-Schneider (Division of Biostatistics) and Nils Becker (Division of Theoretical Systems Biology) for expert advice on data analysis, Joschka Hey (Division of Cancer Epigenomics) for critical advice on ATAC sequencing experiments, Celine Beyersdörffer, Tobias Fischer (both Division of Cellular Immunology) and Oliver Mücke (Division of Cancer Epigenomics) for expert technical assistance, Katja Schmidt and her team (Microbiological Diagnostics Laboratory of the Central Animal Laboratory) for microbiological diagnostics (all DKFZ), the Genomics Unit of the Centre for Genomic Regulation, Barcelona, for providing Illumina sequencing services, Branko Cirovic, Hans Jörg Fehling, Thomas Höfer, Klaus Rajewsky, Axel Roers and Hedda Wardemann for discussions, and all members of the Rodewald laboratory for ongoing support and discussions.

N.D. was initially supported by a fellowship from the Helmholtz Graduate School for Cancer Research. D.B.L. is supported by Deutsche Krebshilfe Grant 70112574. T.H.W. is supported by CRC 130-P11 of the Deutsche Forschungsgemeinschaft. N.D. and H.-R.R. are supported by DKFZ core funding.

## Author contributions

N.D. conceived the study, designed and performed experiments, interpreted results, and wrote the paper, X.W. analyzed molecular data computationally, E.R.C. and B.E. performed experiments, D.B.L. provided support for ChIP sequencing experiments, T.H.W. made autoantibody experiments, Y.B.-N., R.L.K. and S.A.J. provided *Usp22* flox mice, X.W., D.B.L. and T.H.W. edited the paper, and H.-R.R. supervised the study, interpreted results, and wrote the paper.

## Competing interests

The German Cancer Research Center filed an international patent application related to this work. N.D. and H.-R.R. are listed as inventors. The authors have no additional financial interests.

---

**Figure S1.**
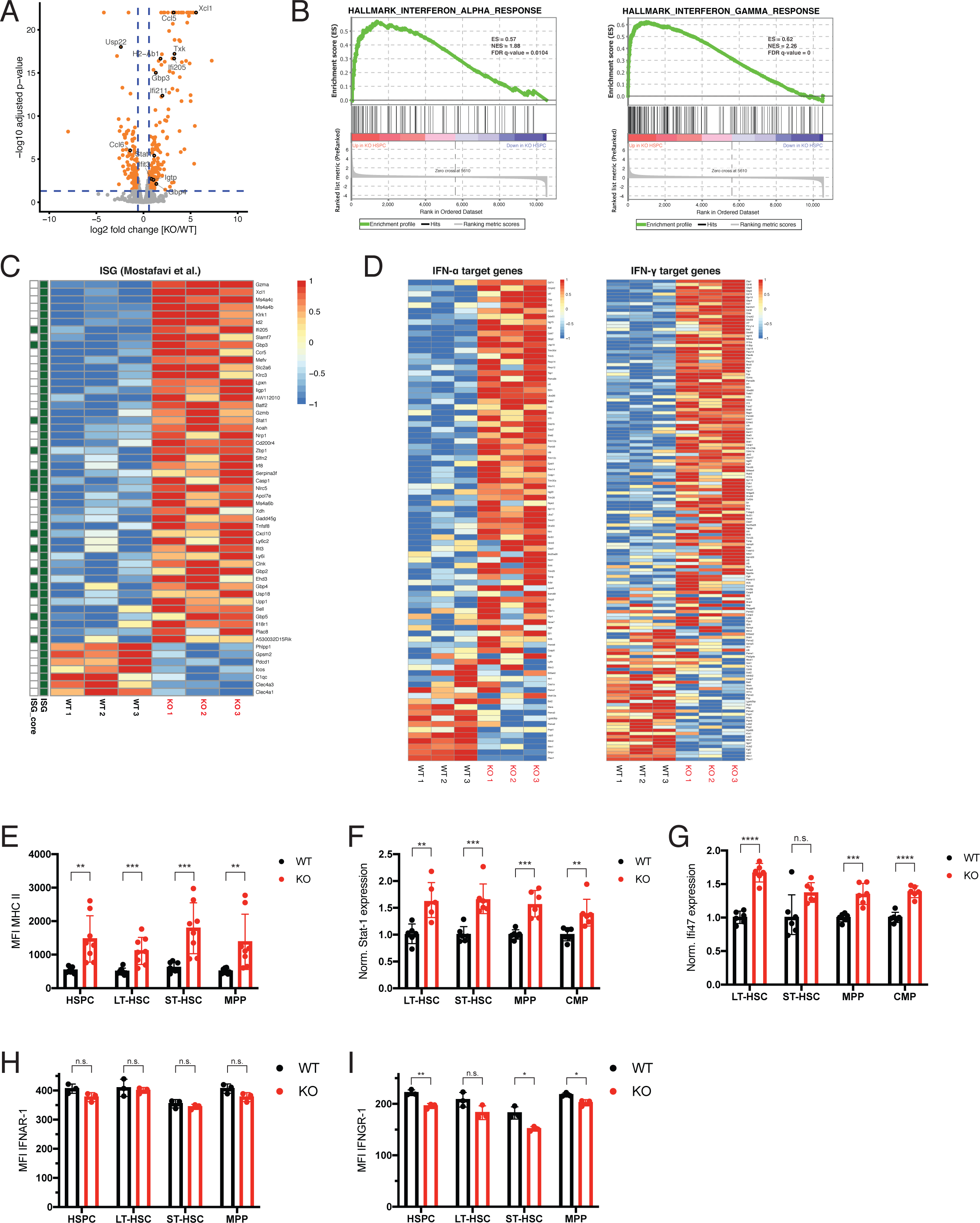
*Usp22* deficiency drives ISG expression. (**A**) Volcano plot showing log2 fold change (FC) in expression and the associated -log10 p-value of individual genes (dots) upon *Usp22* deletion in HSPC based on RNA sequencing. Significantly differentially expressed genes (DEG) (p<0.05, FC *≥* 1.5 or FC *≤* 1/1.5) are highlighted in orange. (**B**) Gene set enrichment analysis (GSEA) plots for IFN-*α* target genes (left panel) and IFN-*γ* target genes (right panel) comparing RNA sequencing data from wild type and *Usp22* KO HSPC. (**C**) Expression levels of all differentially expressed immune cell ISG (Mostafavi et al., 2016) in HSPC isolated from wild type and *Usp22* KO mice. (**D**) Heatmaps of expression of IFN-*α* target genes (left panel) and IFN-*γ* target genes (right panel) in pre-B cells (Fraction C) from wild type and *Usp22* KO mice. (**E**) MHC II median fluorescence intensity (MFI) in wild type and *Usp22* KO stem and progenitor cell populations (n=8). (**F** and **G**) qRT-PCR analysis for *Stat-1* (**F**) and *Ifi47* (**G**) in wild type and *Usp22* KO stem and progenitor cell populations (n=6). (**H** and **I**) Median fluorescence intensity (MFI) of IFNAR-1 (**H**) and IFNGR-1 (**I**) in wild type and *Usp22* KO stem and progenitor cell populations (n=3). Mean ± s.d (**E, H, I**), geometric mean ± geometric s.d. (**F**, **G**). * p<0.05; ** p<0.01; *** p<0.001; **** p<0.0001; n.s., not significant by unpaired, two-tailed Student’s t-test.

**Figure S2.**
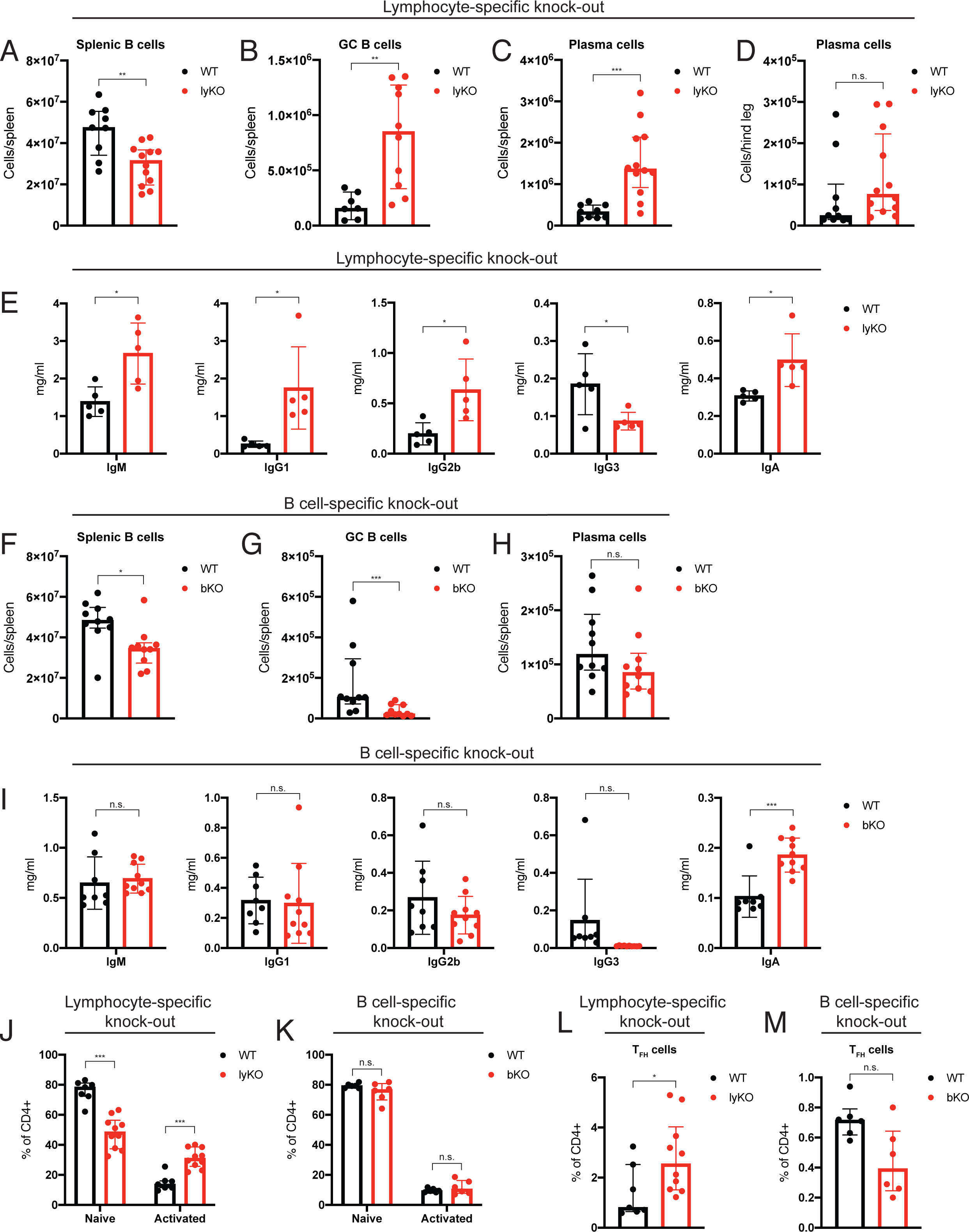
Immunoglobulin hyperproduction upon loss of *Usp22* occurs in a lymphocyte-intrinsic but B cell-extrinsic manner. (**A**-**D**), Absolute numbers of CD19^+^B220^+^ B cells (n=9-12) (**A**), GC B cells (n=7-10) (**B**) and PC (n=9-12) (**C**) in the spleens, and PC in bone marrow (n=10-12) (**D**) of wild type and lymphocyte-specific *Usp22* knock-out (lyKO) mice. (**E**) Serum concentrations of the indicated immunoglobulin isotypes in wild type and *Usp22* lyKO mice (n=5). (**F**-**H**) Absolute numbers of CD19^+^B220^+^ B cells (n=10) (**F**), GC B cells (n=10) (**G**) and PC (n=10) (**H**) in the spleens of wild type and B cell-specific *Usp22* knock-out (bKO) mice. (**I**) Serum concentrations of the indicated immunoglobulin isotypes in wild type and *Usp22* bKO mice (n=8-10). (**J** and **K**) Relative proportions of CD62L^+^CD44^low^ naïve and CD62L^-^CD44^high^ activated CD4^+^ T cells in the spleens of wild type and *Usp22* lyKO mice (n=7-10) (**J**), and in the spleens of wild type and *Usp22* bKO mice (n=6) (**K**). (**L** and **M**) Percentages of PD-1^+^CXCR5^+^ TFH cells within splenic CD4^+^ T cells of wild type and *Usp22* lyKO mice (n=7-10) (**L**), and of wild type and *Usp22* bKO mice (n=6) (**M**). Median ± interquartile range (**A**-**D**, **F**-**H**, **J**-**M**), mean ± s.d. (**E** and **I**). * p<0.05; ** p<0.01; *** p<0.001; **** p<0.0001; n.s., not significant by unpaired, two-tailed Mann-Whitney test (**A**-**D**, **F**-**H**, **J**-**M**) or unpaired, two-tailed Student’s t-test (**E** and **I**).

**Figure S3.**
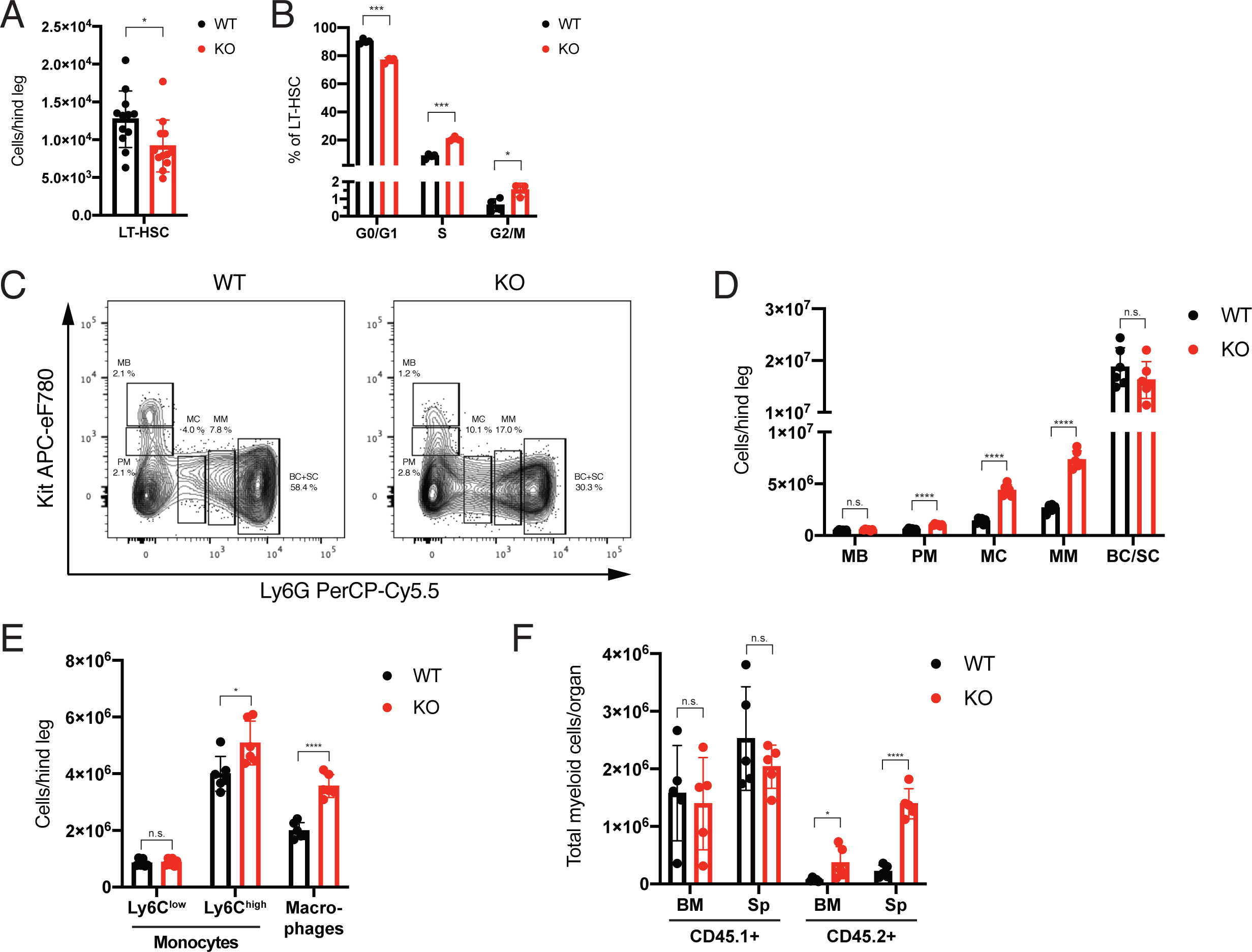
Enhanced myelopoiesis in the absence of *Usp22.* (**A**) Absolute numbers of LT-HSC in the bone marrow of wild type and *Usp22* KO mice (n=12). (**B**) Cell cycle analysis of wild type and *Usp22* KO LT-HSC based on EdU incorporation (n=3-4). (**C**) Expression of Kit and Ly6G resolves stages of myelopoiesis, i.e. myeloblasts (MB), promyelocytes (PM), myelocytes (MC), metamyelocytes (MM), band cells (BC) and segmented cells (SC) in wild type and *Usp22* KO BM cells. Cells were pre-gated on Lineage marker (B220, CD4, CD8 and Ter119)^-^ cells, and using a non-MEP gate. (**D**) Absolute numbers of MB, PM, MC, MM and BC/SC (gated as in **C**) in the bone marrow of wild type and *Usp22* KO mice (n=6). (**E**) Absolute numbers of Ly6C^low^ and Ly6C^high^ monocytes, and macrophages in the bone marrow of wild type and *Usp22* KO mice (n=6). (**F**) Absolute numbers of CD45.1^+^ and CD45.2^+^ myeloid cells in the bone marrow and spleens of mice transplanted with wild type CMP, or *Usp22* KO CMP (both CD45.2^+^) together with wild type (CD45.1^+^) total bone marrow competitor cells 10 days after transplantation (n=5). Mean ± s.d. * p<0.05; ** p<0.01; *** p<0.001; **** p<0.0001; n.s., not significant by unpaired, two-tailed Student’s t-test.

**Figure S4.**
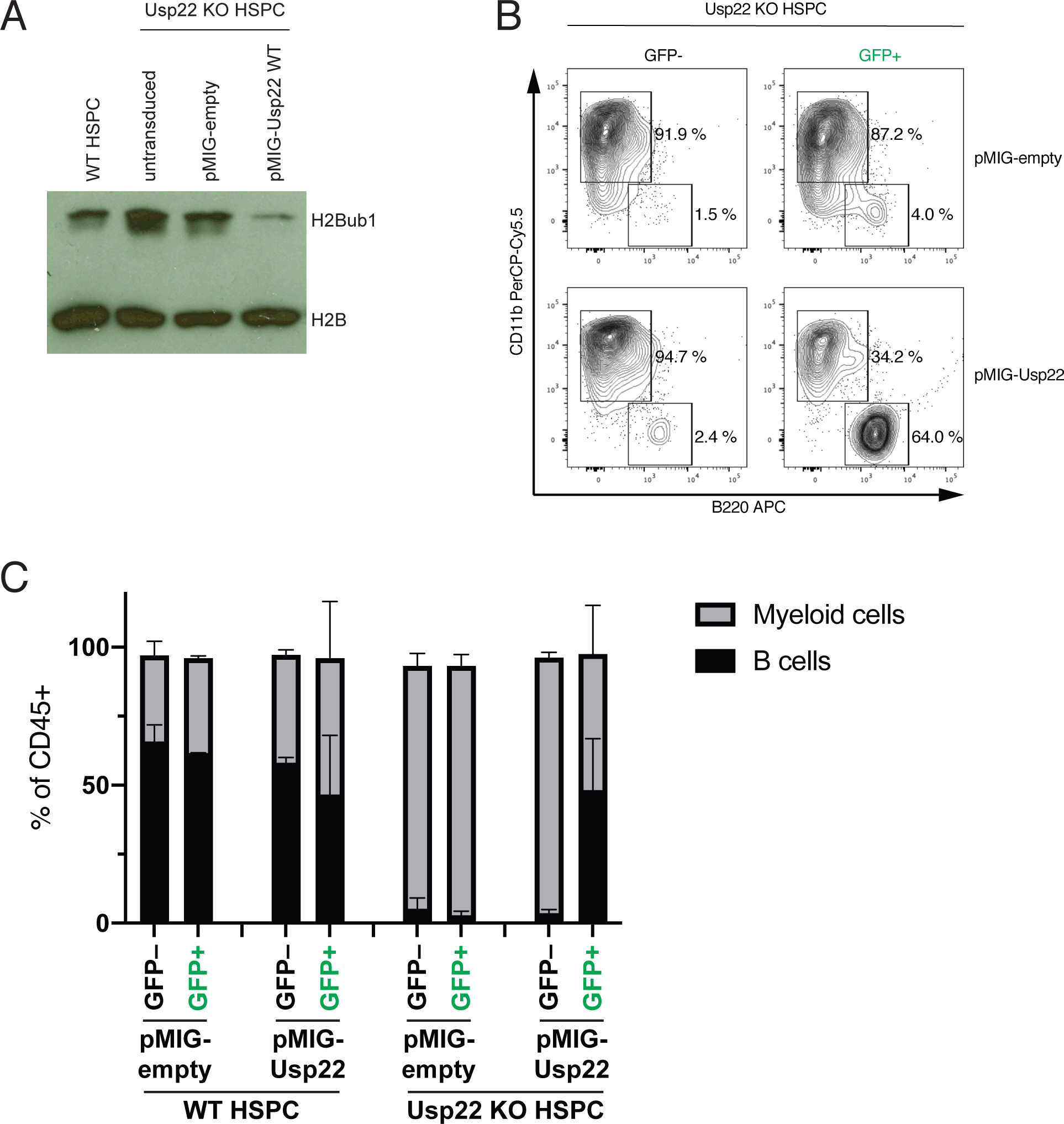
Re-expression of *Usp22* in *Usp22* KO HSPC reverts the myeloid bias in vitro. (**A**) Western Blot for H2B and H2Bub1 at day 4 post transduction of wild type and *Usp22* KO HSPC with empty pMSCV-IRES-GFP (pMIG) or pMIG-*Usp22*. (**B**) CD11b and B220 expression in cells differentiated in vitro for 1.5 weeks from *Usp22* KO HSPC transduced with the indicated vectors. Transduced cells can be identified based on the expression of GFP. (**C**) Relative proportions of B220^+^ B cells and CD11b^+^ myeloid cells (gated as in **B**) within cells differentiated for 1.5 weeks in vitro from wild type and *Usp22* KO HSPC transduced with the indicated gamma-retroviral expression vectors (n=3). The partial myeloid cell production in these experiments also from wild type progenitors presumably results from the cellular pre-expansion conditions required for viral transduction prior to the differentiation assay. Mean ± s.d.

**Figure S5.**
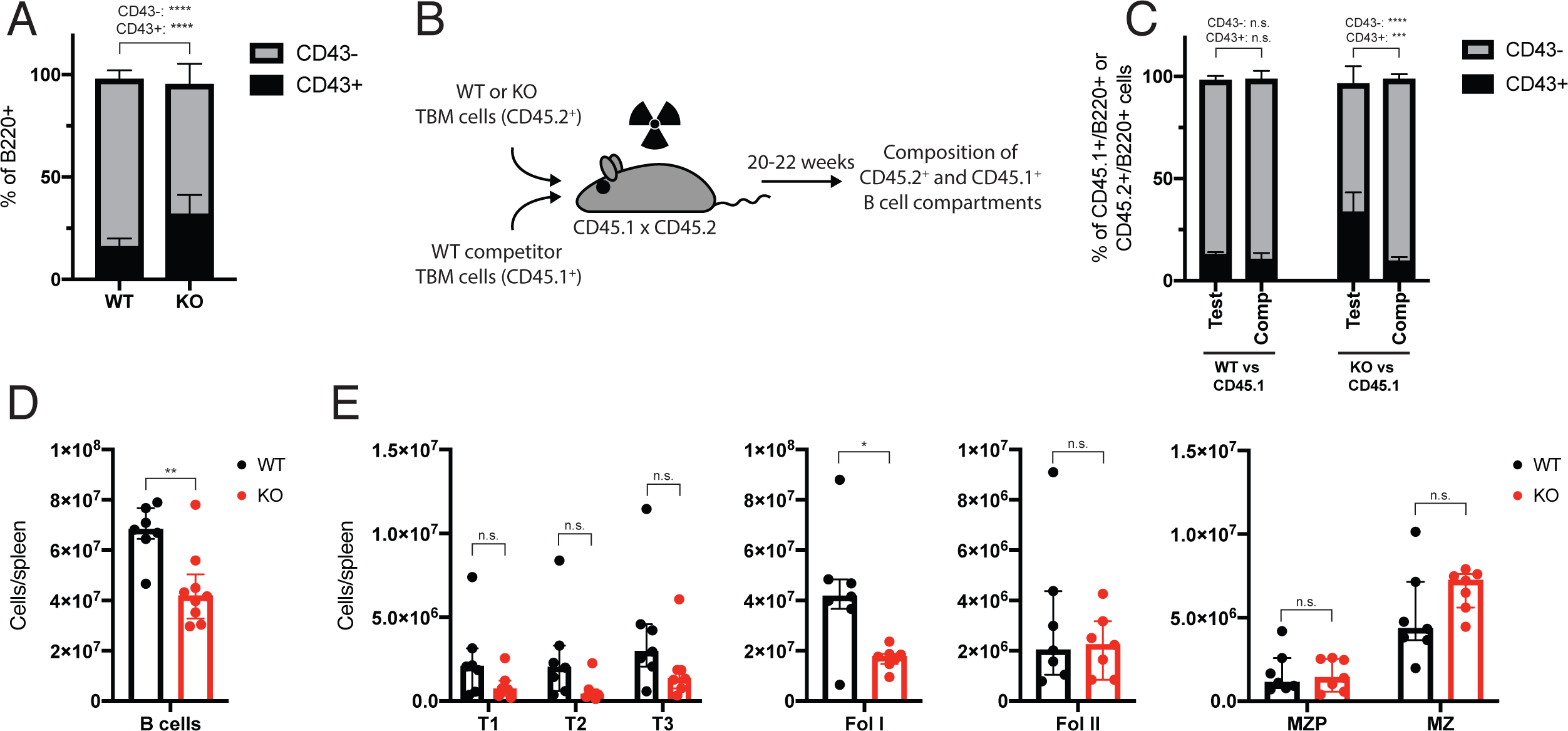
Impaired B cell development in *Usp22* KO mice. (**A**) Relative proportions of early CD43^+^ and late CD43^-^ B cell progenitors within the B220^+^ population of wild type and *Usp22* KO mice (n=12). (**B**) Experimental setup for the generation of mixed bone marrow chimeras. (**C**) Relative proportions of early CD43^+^ and late CD43^-^ B cell progenitors within the B220^+^ population of CD45.2^+^ test and CD45.1^+^ competitor cells in mixed bone marrow chimeras (n=6). (**D**) Absolute numbers of CD19^+^B220^+^ splenic B cells in wild type and *Usp22* KO mice (n=7-9). (**E**) Absolute numbers of the indicated splenic B cell subsets in wild type and *Usp22* KO mice (n=7). Mean ± s.d. (**A**, **C**), median ± interquartile range (**D**, **E**). * p<0.05, ** p<0.01, *** p<0.001, **** p<0.0001 by unpaired, two-tailed Student’s t-test (**A**, **C**) or unpaired, two-tailed Mann-Whitney test (**D**, **E**).

**Figure S6.**
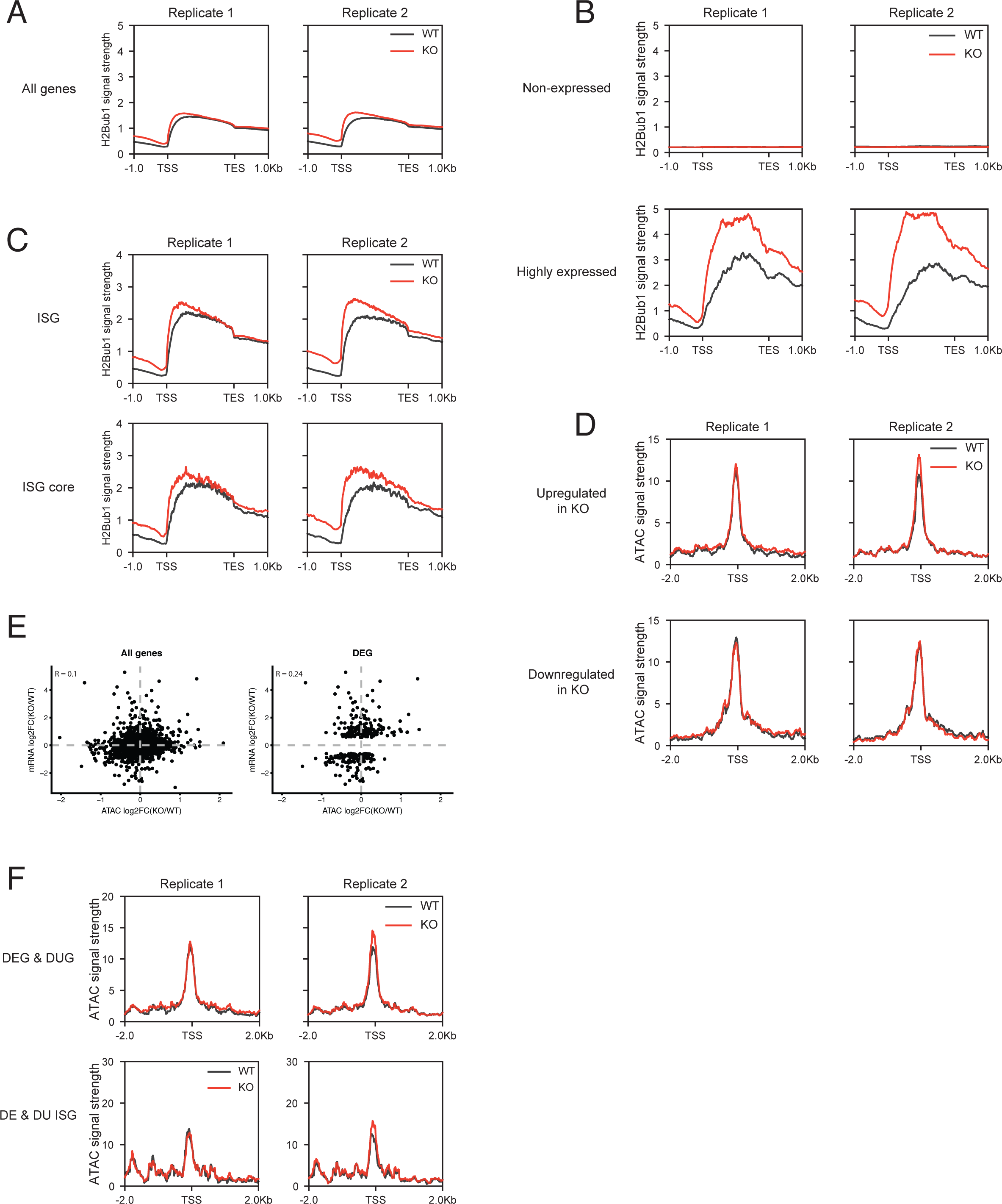
*Usp22* KO cells show increased H2Bub1 levels at ISG loci. (**A**-**C**) Average H2Bub1 ChIP sequencing signal in wild type and *Usp22* KO HSPC across gene bodies of all genes (**A**), of non-expressed genes (**B**, top panels), of genes highly expressed in HSPC (**B**, bottom panels), of all immune cell ISG (Mostafavi et al., 2016) (**C**, top panels), and of the immune cell core ISG (Mostafavi et al., 2016) (**C**, bottom panels). (**D**) Average ATAC sequencing signal around the TSS of genes upregulated in *Usp22* KO HSPC (top panels), and of genes downregulated in *Usp22* KO HSPC (bottom panels) comparing wild type and *Usp22* KO HSPC. (**E**) Correlation between changes in TSS accessibility (measured by ATAC sequencing) and changes in mRNA expression comparing *Usp22* KO and wild type HSPC. Each dot represents an individual gene, left panel shows all genes, and right panel only genes differentially expressed between wild type and *Usp22* KO HSPC. (**F**) Average ATAC sequencing signal around the TSS of differentially expressed and ubiquitinated genes (top panel), and differentially expressed (DE) and differentially ubiquitinated (DU) ISG (bottom panel) in wild type and *Usp22* KO HSPC. TSS = Transcription start site; TES = Transcription end site.

**Table S1.** Extended survey for the presence of pathogens in wild type and Usp22-deficient mice reveals no signs of infections. Shown is the fraction of mice positively tested for the indicated pathogens (n=4).

| Methodology | Infectious agent | WT<br>(positive/total) | KO<br>(positive/total) |
| --- | --- | --- | --- |
| Microbiology | Bacterial infectious agents | 0/4 | 0/4 |
| Molecular biology | Demodex muris | 0/4 | 0/4 |
|  | Helicobacter spp. | 0/4 | 0/4 |
|  | Mites | 0/4 | 0/4 |
|  | Mouse kidney parvovirus | 0/4 | 0/4 |
|  | Pneumocystis murina | 0/4 | 0/4 |
| Parasitology<br>(Microscopic) | Ectoparasites | 0/4 | 0/4 |
|  | Endoparasites | 0/4 | 0/4 |
| Serology<br>(Bacteria) | Clostridium piliforme | 0/4 | 0/4 |
|  | Muribacter muris | 0/4 | 0/4 |
|  | Mycoplasma pulmonis | 0/4 | 0/4 |
|  | Rodentibacter spp. | 0/4 | 0/4 |
|  | Streptobacillus moniliformis | 0/4 | 0/4 |
| Serology<br>(Viruses) | Ectromelia virus | 0/4 | 0/4 |
|  | Encephalomyocarditis virus | 0/4 | 0/4 |
|  | Hantaan virus | 0/4 | 0/4 |
|  | K virus | 0/4 | 0/4 |
|  | Lactate dehydrogenase elevating virus | 0/4 | 0/4 |
|  | Lymphocytic choriomeningitis virus | 0/4 | 0/4 |
|  | Mouse adenovirus FL | 0/4 | 0/4 |
|  | Mouse adenovirus K87 | 0/4 | 0/4 |
|  | Mouse hepatitis virus | 0/4 | 0/4 |
|  | Mouse minute virus | 0/4 | 0/4 |
|  | Mouse parvovirus | 0/4 | 0/4 |
|  | Mouse rotavirus (EDIM) | 0/4 | 0/4 |
|  | Murine astrovirus | 0/4 | 0/4 |
|  | Murine cytomegalovirus | 0/4 | 0/4 |
|  | Murine norovirus | 0/4 | 0/4 |
|  | Parvoviruses | 0/4 | 0/4 |
|  | Polyoma virus | 0/4 | 0/4 |
|  | Puumala virus | 0/4 | 0/4 |
|  | Reovirus 3 | 0/4 | 0/4 |
|  | Sendai virus | 0/4 | 0/4 |
|  | Theiler's GD VII virus | 0/4 | 0/4 |

**Table S2.**
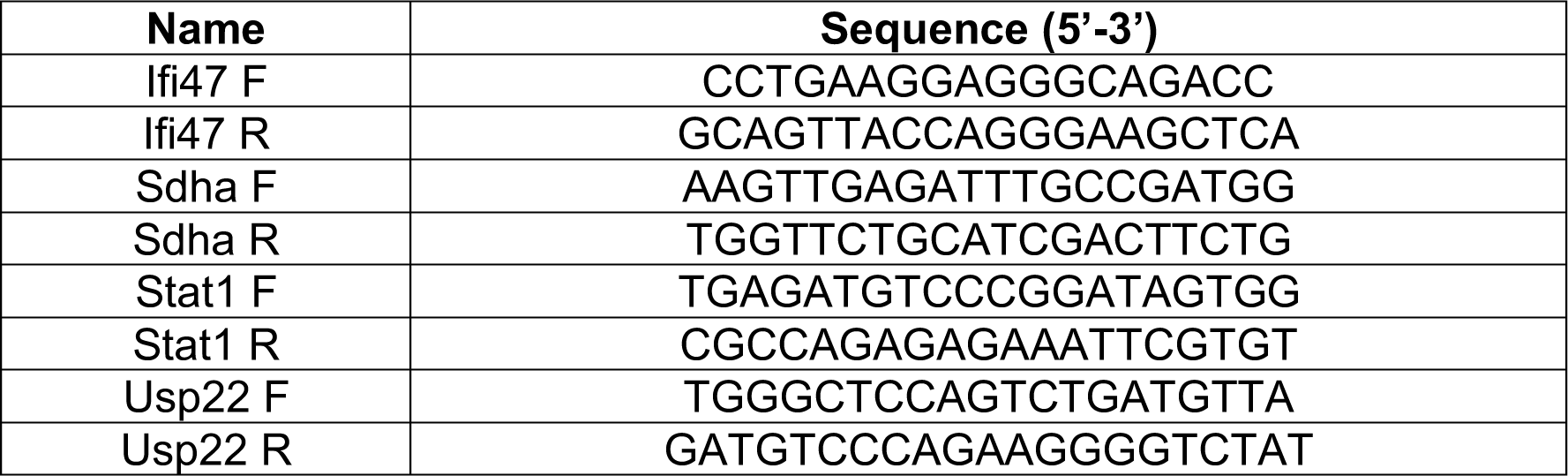
Primers used for qRT-PCR.

**Table S3.**
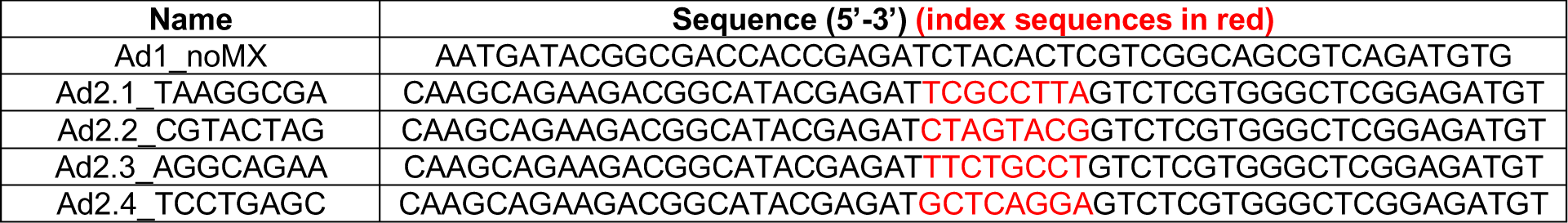
Indexing primers used for the preparation of ChIP and ATAC sequencing libraries.

## Notes

### Summary of Updates

Funding sources have been corrected.

## References

1. Anders, S., P.T. Pyl, and W. Huber. 2015. HTSeq--a Python framework to work with high-throughput sequencing data. Bioinformatics. 31:166–169. doi:10.1093/bioinformatics/btu638.

2. Baldridge, M.T., K.Y. King, N.C. Boles, D.C. Weksberg, and M.A. Goodell. 2010. Quiescent haematopoietic stem cells are activated by IFN-γ in response to chronic infection. Nature. 465:793–797. doi:10.1038/nature09135.

3. Buechler, M.B., T.H. Teal, K.B. Elkon, and J.A. Hamerman. 2013. Cutting Edge: Type I IFN Drives Emergency Myelopoiesis and Peripheral Myeloid Expansion during Chronic TLR7 Signaling. J. Immunol. 190:886–891. doi:10.4049/jimmunol.1202739.

4. Buenrostro, J.D., B. Wu, H.Y. Chang, and W.J. Greenleaf. 2015. ATAC-seq: A Method for Assaying Chromatin Accessibility Genome-Wide. Curr. Protoc. Mol. Biol. 109:1–9. doi:10.1002/0471142727.mb2129s109.

5. Cai, Z., M.-X. Zhang, Z. Tang, Q. Zhang, J. Ye, T.-C. Xiong, Z.-D. Zhang, and B. Zhong. 2020. USP22 promotes IRF3 nuclear translocation and antiviral responses by deubiquitinating the importin protein KPNA2. J. Exp. Med. 217:e20191174. doi:10.1084/jem.20191174.

6. Cho, S.K., T.D. Webber, J.R. Carlyle, T. Nakano, S.M. Lewis, and J.C. Zúñiga-Pflücker. 1999. Functional characterization of B lymphocytes generated in vitro from embryonic stem cells. Proc. Natl. Acad. Sci. USA. 96:9797–9802. doi:10.1073/pnas.96.17.9797.

7. Corces, M.R., A.E. Trevino, E.G. Hamilton, P.G. Greenside, N.A. Sinnott-Armstrong, S. Vesuna, A.T. Satpathy, A.J. Rubin, K.S. Montine, B. Wu, A. Kathiria, S.W. Cho, M.R. Mumbach, A.C. Carter, M. Kasowski, L.A. Orloff, V.I. Risca, A. Kundaje, P.A. Khavari, T.J. Montine, W.J. Greenleaf, and H.Y. Chang. 2017. An improved ATAC-seq protocol reduces background and enables interrogation of frozen tissues. Nat. Methods. 14:959– 962. doi:10.1038/nmeth.4396.

8. Cortez, J.T., E. Montauti, E. Shifrut, J. Gatchalian, Y. Zhang, O. Shaked, Y. Xu, T.L. Roth, D.R. Simeonov, Y. Zhang, S. Chen, Z. Li, J.M. Woo, J. Ho, I.A. Vogel, G.Y. Prator, Bin Zhang, Y. Lee, Z. Sun, I. Ifergan, F. Van Gool, D.C. Hargreaves, J.A. Bluestone, A. Marson, and D. Fang. 2020. CRISPR screen in regulatory T cells reveals modulators of Foxp3. Nature. 582:416–420. doi:10.1038/s41586-020-2246-4.

9. Crow, M.K., M. Olferiev, and K.A. Kirou. 2019. Type I Interferons in Autoimmune Disease. Annu. Rev. Pathol. Mech. Dis. 14:369–393. doi:10.1146/annurev-pathol-020117-043952.

10. de Boer, J., A. Williams, G. Skavdis, N. Harker, M. Coles, M. Tolaini, T. Norton, K. Williams, K. Roderick, A.J. Potocnik, and D. Kioussis. 2003. Transgenic mice with hematopoietic and lymphoid specific expression of Cre. Eur. J. Immunol. 33:314–325. doi:10.1002/immu.200310005.

11. Dobin, A., C.A. Davis, F. Schlesinger, J. Drenkow, C. Zaleski, S. Jha, P. Batut, M. Chaisson, and T.R. Gingeras. 2012. STAR: ultrafast universal RNA-seq aligner. Bioinformatics. 29:15–21. doi:10.1093/bioinformatics/bts635.

12. Dodt, M., J.T. Roehr, R. Ahmed, and C. Dieterich. 2012. FLEXBAR-Flexible Barcode and Adapter Processing for Next-Generation Sequencing Platforms. Biology (Basel). 1:895– 905. doi:10.3390/biology1030895.

13. Essers, M.A.G., S. Offner, W.E. Blanco-Bose, Z. Waibler, U. Kalinke, M.A. Duchosal, and A. Trumpp. 2009. IFNα activates dormant haematopoietic stem cells in vivo. Nature. 458:904–908. doi:10.1038/nature07815.

14. Fonseca, G.J., G. Thillainadesan, A.F. Yousef, J.N. Ablack, K.L. Mossman, J. Torchia, and J.S. Mymryk. 2012. Adenovirus Evasion of Interferon-Mediated Innate Immunity by Direct Antagonism of a Cellular Histone Posttranslational Modification. Cell Host Microbe. 11:597–606. doi:10.1016/j.chom.2012.05.005.

15. Grawunder, U., F. Melchers, and A. Rolink. 1993. Interferon-gamma arrests proliferation and causes apoptosis in stromal cell/interleukin-7-dependent normal murine pre-B cell lines and clones in vitro, but does not induce differentiation to surface immunoglobulin-positive B cells. Eur. J. Immunol. 23:544–551. doi:10.1002/eji.1830230237.

16. Hardy, R.R., C.E. Carmack, S.A. Shinton, J.D. Kemp, and K. Hayakawa. 1991. Resolution and characterization of pro-B and pre-pro-B cell stages in normal mouse bone marrow. J. Exp. Med. 173:1213–1225. doi:10.1084/jem.173.5.1213.

17. Ivashkiv, L.B., and L.T. Donlin. 2013. Regulation of type I interferon responses. Nat. Rev. Immunol. 14:36–49. doi:10.1038/nri3581.

18. Jego, G., A.K. Palucka, J.-P. Blanck, C. Chalouni, V. Pascual, and J. Banchereau. 2003. Plasmacytoid Dendritic Cells Induce Plasma Cell Differentiation through Type I Interferon and Interleukin 6. Immunity. 19:225–234. doi:10.1016/S1074-7613(03)00208-5.

19. Kim, D., G. Pertea, C. Trapnell, H. Pimentel, R. Kelley, and S.L. Salzberg. 2013. TopHat2: accurate alignment of transcriptomes in the presence of insertions, deletions and gene fusions. Genome Biol. 14:R36. doi:10.1186/gb-2013-14-4-r36.

20. Langmead, B., and S.L. Salzberg. 2012. Fast gapped-read alignment with Bowtie 2. Nat. Methods. 9:357–359. doi:10.1038/nmeth.1923.

21. Li, C., T. Irrazabal, C.C. So, M. Berru, L. Du, E. Lam, A.K. Ling, J.L. Gommerman, Q. Pan-Hammarström, and A. Martin. 2018. The H2B deubiquitinase Usp22 promotes antibody class switch recombination by facilitating non-homologous end joining. Nat. Commun. 9:1–12. doi:10.1038/s41467-018-03455-x.

22. Liberzon, A., C. Birger, H. Thorvaldsdóttir, M. Ghandi, J.P. Mesirov, and P. Tamayo. 2015. The Molecular Signatures Database (MSigDB) hallmark gene set collection. Cell Syst. 1:417–425. doi:10.1016/j.cels.2015.12.004.

23. Lin, Q., C. Dong, and M.D. Cooper. 1998. Impairment of T and B cell development by treatment with a type I interferon. J. Exp. Med. 187:79–87. doi:10.1084/jem.187.1.79.

24. Lin, Z., H. Yang, Q. Kong, J. Li, S.-M. Lee, B. Gao, H. Dong, J. Wei, J. Song, D.D. Zhang, and D. Fang. 2012. USP22 Antagonizes p53 Transcriptional Activation by Deubiquitinating Sirt1 to Suppress Cell Apoptosis and Is Required for Mouse Embryonic Development. Mol. Cell. 46:484–494. doi:10.1016/j.molcel.2012.03.024.

25. Love, M.I., W. Huber, and S. Anders. 2014. Moderated estimation of fold change and dispersion for RNA-seq data with DESeq2. Genome Biol. 15:31–54. doi:10.1186/s13059-014-0550-8.

26. Melo-Cardenas, J., Y. Xu, J. Wei, C. Tan, S. Kong, B. Gao, E. Montauti, G. Kirsammer, J.D. Licht, J. Yu, P. Ji, J.D. Crispino, and D. Fang. 2018. USP22 deficiency leads to myeloid leukemia upon oncogenic Kras activation through a PU.1-dependent mechanism. Blood. 132:423–434. doi:10.1182/blood-2017-10-811760.

27. Minsky, N., E. Shema, Y. Field, M. Schuster, E. Segal, and M. Oren. 2008. Monoubiquitinated H2B is associated with the transcribed region of highly expressed genes in human cells. Nat. Cell Biol. 10:483–488. doi:10.1038/ncb1712.

28. Mostafavi, S., H. Yoshida, D. Moodley, H. LeBoité, K. Rothamel, T. Raj, C.J. Ye, N. Chevrier, S.-Y. Zhang, T. Feng, M. Lee, J.-L. Casanova, J.D. Clark, M. Hegen, J.-B. Telliez, N. Hacohen, P.L. De Jager, A. Regev, D. Mathis, C. Benoist, and T.I.G.P. Consortium. 2016. Parsing the Interferon Transcriptional Network and Its Disease Associations. Cell. 164:564–578. doi:10.1016/j.cell.2015.12.032.

29. Nakano, T., H. Kodama, and T. Honjo. 1994. Generation of lymphohematopoietic cells from embryonic stem cells in culture. Science. 265:1098–1101. doi:10.1126/science.8066449.

30. Pavri, R., B. Zhu, G. Li, P. Trojer, S. Mandal, A. Shilatifard, and D. Reinberg. 2006. Histone H2B monoubiquitination functions cooperatively with FACT to regulate elongation by RNA polymerase II. Cell. 125:703–717. doi:10.1016/j.cell.2006.04.029.

31. Prenzel, T., Y. Begus-Nahrmann, F. Kramer, M. Hennion, C. Hsu, T. Gorsler, C. Hintermair, D. Eick, E. Kremmer, M. Simons, T. Beissbarth, and S.A. Johnsen. 2011. Estrogen-Dependent Gene Transcription in Human Breast Cancer Cells Relies upon Proteasome-Dependent Monoubiquitination of Histone H2B. Cancer Res. 71:5739–5753. doi:10.1158/0008-5472.CAN-11-1896.

32. Ramírez, F., D.P. Ryan, B. Grüning, V. Bhardwaj, F. Kilpert, A.S. Richter, S. Heyne, F. Dündar, and T. Manke. 2016. deepTools2: a next generation web server for deep-sequencing data analysis. Nucleic Acids Res. 44:W160–165. doi:10.1093/nar/gkw257.

33. Rickert, R.C., J. Roes, and K. Rajewsky. 1997. B lymphocyte-specific, Cre-mediated mutagenesis in mice. Nucleic Acids Res. 25:1317–1318. doi:10.1093/nar/25.6.1317.

34. Sarkar, D., E.S. Park, and P.B. Fisher. 2006. Defining the mechanism by which IFN-beta dowregulates c-myc expression in human melanoma cells: pivotal role for human polynucleotide phosphorylase (hPNPaseold-35). Cell Death Differ. 13:1541–1553. doi:10.1038/sj.cdd.4401829.

35. Schlenner, S.M., V. Madan, K. Busch, A. Tietz, C. LAufle, C. Costa, C. Blum, H.J. Fehling, and H.-R. Rodewald. 2010. Fate Mapping Reveals Separate Origins of T Cells and Myeloid Lineages in the Thymus. Immunity. 32:426–436. doi:10.1016/j.immuni.2010.03.005.

36. Schmidl, C., A.F. Rendeiro, N.C. Sheffield, and C. Bock. 2015. ChIPmentation: fast, robust, low-input ChIP-seq for histones and transcription factors. Nat. Methods. 12:963– 965. doi:10.1038/nmeth.3542.

37. Schneider, W.M., M.D. Chevillotte, and C.M. Rice. 2014. Interferon-Stimulated Genes: A Complex Web of Host Defenses. Annu. Rev. Immunol. 32:513–545. doi:10.1146/annurev-immunol-032713-120231.

38. Shema, E., I. Tirosh, Y. Aylon, J. Huang, C. Ye, N. Moskovits, N. Raver-Shapira, N. Minsky, J. Pirngruber, G. Tarcic, P. Hublarova, L. Moyal, M. Gana-Weisz, Y. Shiloh, Y. Yarden, S.A. Johnsen, B. Vojtesek, S.L. Berger, and M. Oren. 2008. The histone H2B-specific ubiquitin ligase RNF20/hBRE1 acts as a putative tumor suppressor through selective regulation of gene expression. Genes Dev. 22:2664–2676. doi:10.1101/gad.1703008.

39. Shukla, A., and S.R. Bhaumik. 2007. H2B-K123 ubiquitination stimulates RNAPII elongation independent of H3-K4 methylation. Biochem. Biophys. Res. Commun. 359:214–220. doi:10.1016/j.bbrc.2007.05.105.

40. Subramanian, A., P. Tamayo, V.K. Mootha, S. Mukherjee, B.L. Ebert, M.A. Gillette, A. Paulovich, S.L. Pomeroy, T.R. Golub, E.S. Lander, and J.P. Mesirov. 2005. Gene set enrichment analysis: a knowledge-based approach for interpreting genome-wide expression profiles. Proc. Natl. Acad. Sci. USA. 102:15545–15550. doi:10.1073/pnas.0506580102.

41. Vallespinós, M., D. Fernández, L. Rodríguez, J. Alvaro-Blanco, E. Baena, M. Ortiz, D. Dukovska, D. Martínez, A. Rojas, M.R. Campanero, and I. Moreno de Alborán. 2011. B Lymphocyte Commitment Program Is Driven by the Proto-Oncogene c-myc. J.Immunol. 186:6726–6736. doi:10.4049/jimmunol.1002753.

42. Xie, W., S. Nagarajan, S.J. Baumgart, R.L. Kosinsky, Z. Najafova, V. Kari, M. Hennion, D. Indenbirken, S. Bonn, A. Grundhoff, F. Wegwitz, A. Mansouri, and S.A. Johnsen. 2017. RNF40 regulates gene expression in an epigenetic context-dependent manner. Genome Biol. 18:32. doi:10.1186/s13059-017-1159-5.

43. Young, M.D., M.J. Wakefield, G.K. Smyth, and A. Oshlack. 2010. Gene ontology analysis for RNA-seq: accounting for selection bias. Genome Biol. 11:R14. doi:10.1186/gb-2010-11-2-r14.

44. Zhang, F., and X. Yu. 2011. WAC, a Functional Partner of RNF20/40, Regulates Histone H2B Ubiquitination and Gene Transcription. Mol. Cell. 41:384–397. doi:10.1016/j.molcel.2011.01.024.

45. Zhang, X.-Y., M. Varthi, S.M. Sykes, C. Phillips, C. Warzecha, W. Zhu, A. Wyce, A.W. Thorne, S.L. Berger, and S.B. McMahon. 2008a. The Putative Cancer Stem Cell Marker USP22 Is a Subunit of the Human SAGA Complex Required for Activated Transcription and Cell-Cycle Progression. Mol. Cell. 29:102–111. doi:10.1016/j.molcel.2007.12.015.

46. Zhang, Y., T. Liu, C.A. Meyer, J. Eeckhoute, D.S. Johnson, B.E. Bernstein, C. Nusbaum, R.M. Myers, M. Brown, W. Li, and X.S. Liu. 2008b. Model-based analysis of ChIP-Seq (MACS). Genome Biol. 9:R137. doi:10.1186/gb-2008-9-9-r137.

47. Zhao, Y., G. Lang, S. Ito, J. Bonnet, E. Metzger, S. Sawatsubashi, E. Suzuki, X. Le Guezennec, H.G. Stunnenberg, A. Krasnov, S.G. Georgieva, R. Schüle, K.-I. Takeyama, S. Kato, L. Tora, and D. Devys. 2008. A TFTC/STAGA Module Mediates Histone H2A and H2B Deubiquitination, Coactivates Nuclear Receptors, and Counteracts Heterochromatin Silencing. Mol. Cell. 29:92–101. doi:10.1016/j.molcel.2007.12.011.

48. Zhu, B., Y. Zheng, A.-D. Pham, S.S. Mandal, H. Erdjument-Bromage, P. Tempst, and D. Reinberg. 2005. Monoubiquitination of Human Histone H2B: The Factors Involved and Their Roles in HOX Gene Regulation. Mol. Cell. 20:601–611. doi:10.1016/j.molcel.2005.09.025.

